# Cryo-ET reveals the *in situ* architecture of the polar tube invasion apparatus from microsporidian parasites

**DOI:** 10.1101/2024.07.13.603322

**Authors:** Mahrukh Usmani, Nicolas Coudray, Margot Riggi, Rishwanth Raghu, Harshita Ramchandani, Daija Bobe, Mykhailo Kopylov, Ellen D. Zhong, Janet H. Iwasa, Damian C. Ekiert, Gira Bhabha

**Affiliations:** Department of Biology, Johns Hopkins University, Baltimore, MD, USA; Vilcek Institute of Graduate Biomedical Sciences, NYU School of Medicine, New York, NY, USA; Applied Bioinformatics Laboratories, Department of Medicine, NYU School of Medicine, New York, NY, USA; Department of Biochemistry, University of Utah, Salt Lake City, UT, USA; Department of Computer Science, Princeton University, Princeton, NJ, USA; Simons Electron Microscopy Center, New York Structural Biology Center, New York, NY, USA

**Author notes:** equal contribution (Mahrukh Usmani, Nicolas Coudray). corresponding author. (G.B.); (D.C.E.).

## Abstract

Microsporidia are divergent fungal pathogens that employ a harpoon-like apparatus called the polar tube (PT) to invade host cells. The PT architecture and its association with neighboring organelles remain poorly understood. Here, we use cryo-electron tomography to investigate the structural cell biology of the PT in dormant spores from the human-infecting microsporidian species, *Encephalitozoon intestinalis*. Segmentation and subtomogram averaging of the PT reveal at least four layers: two protein-based layers surrounded by a membrane, and filled with a dense core. Regularly spaced protein filaments form the structural skeleton of the PT. Combining cryo-electron tomography with cellular modeling, we propose a model for the 3-dimensional organization of the polaroplast, an organelle that is continuous with the membrane layer that envelops the PT. Our results reveal the ultrastructure of the microsporidian invasion apparatus *in situ*, laying the foundation for understanding infection mechanisms.

## Introduction

To initiate infection, intracellular eukaryotic parasites have evolved a wide-range of invasion organelles that mediate parasite attachment and entry into host cells. For example, in apicomplexan parasites, such as *Plasmodium falciparum, Toxoplasma gondii*, and *Cryptosporidium parvum*, specialized secretory organelles called rhoptries export numerous proteins to the interface between the parasite and host cells, which alter host cell physiology and facilitate parasite invasion^1,2^. In some fungal pathogens of plants, such as *Magnaporthe oryzae*, a “pressing” organelle called the appressorium generates very high mechanical force at the tip of a narrow cellular projection that is used to punch a hole through the rigid cell wall of the plant cell, allowing a fungal hypha to gain entry to the host cell interior^3^. In microsporidia, which are an early-diverging group of fungi^4,5^, a unique harpoon-like ballistic organelle called the polar tube (PT) mediates invasion of host cells^6–8^. While the microsporidian PT was discovered more than 100 years ago, how the PT and its associated organelles interact to mediate invasion remains poorly understood relative to the invasion mechanisms and organelles of other eukaryotic pathogens.

Over 1,700 species of microsporidia are known^9^ that can infect a wide range of animal hosts. About 15 microsporidian species cause disease in humans^10–12^, while other species infect economically and ecologically important animals including livestock^13–15^, insects^16–18^, and aquatic animals^19–22^. In immunocompromised patients, opportunistic microsporidian infections can cause fatal illnesses^23– 25^. Microsporidia are transmitted between hosts in the form of dormant, environmentally-resistant spores. As obligate intracellular pathogens, microsporidia are dependent on their hosts to replicate and complete their life cycle^26^. Central to the microsporidian infection process is the PT, which mediates invasion of a host cell^7,27–29^. In a dormant parasite, the PT is coiled within the spore as a right-handed helix (pre-fired state)^27^, and is associated with two organelles called the polaroplast and anchoring disk, which have been proposed to function together with the PT during the firing and invasion process^30,31^. When triggered, the PT, many times the length of the spore itself, fires rapidly from the spore on the millisecond timescale, and undergoes a large-scale conformational change to transform into a long, linear tube (post-fired state) (**Fig. 1a)**. One end of the PT remains tightly associated with the spore body via the anchoring disk, while the other end of the PT is thought to pierce the host cell membrane, and transport infectious cargo (“sporoplasm”) into the host^27,32^. During this process, it has been proposed that the polaroplast, an elaborate membranous organelle at the anterior end of the dormant spore, provides a reservoir of membrane to facilitate parasite translocation through the PT^33–35^. How the polaroplast is organized in 3D has been challenging to establish, due to its intricate combination of membrane stacks and folds that appear quite variable both within and between species^36–41^. Given the well-defined shape of the PT when viewed by light microscopy, and its apparent mechanical properties that allow substantial force generation to penetrate a host cell, it seems likely that a protein skeleton establishes and maintains the overall architecture of the PT. Several polar tube proteins (PTPs) have been identified that localize to the PT^42–47^, though it remains unclear which proteins are the major structural components. Conventional TEM of spores as well as more recent cryo-ET studies of the PT in its post-fired state have yielded some initial insights into concentric layers of density in the tube wall, as well as the cargo passing through the tube^44,48–50^. However, the complex 3D organization and architecture of the nanometer-to-micron PT machine remains mysterious.

**Figure 1.**
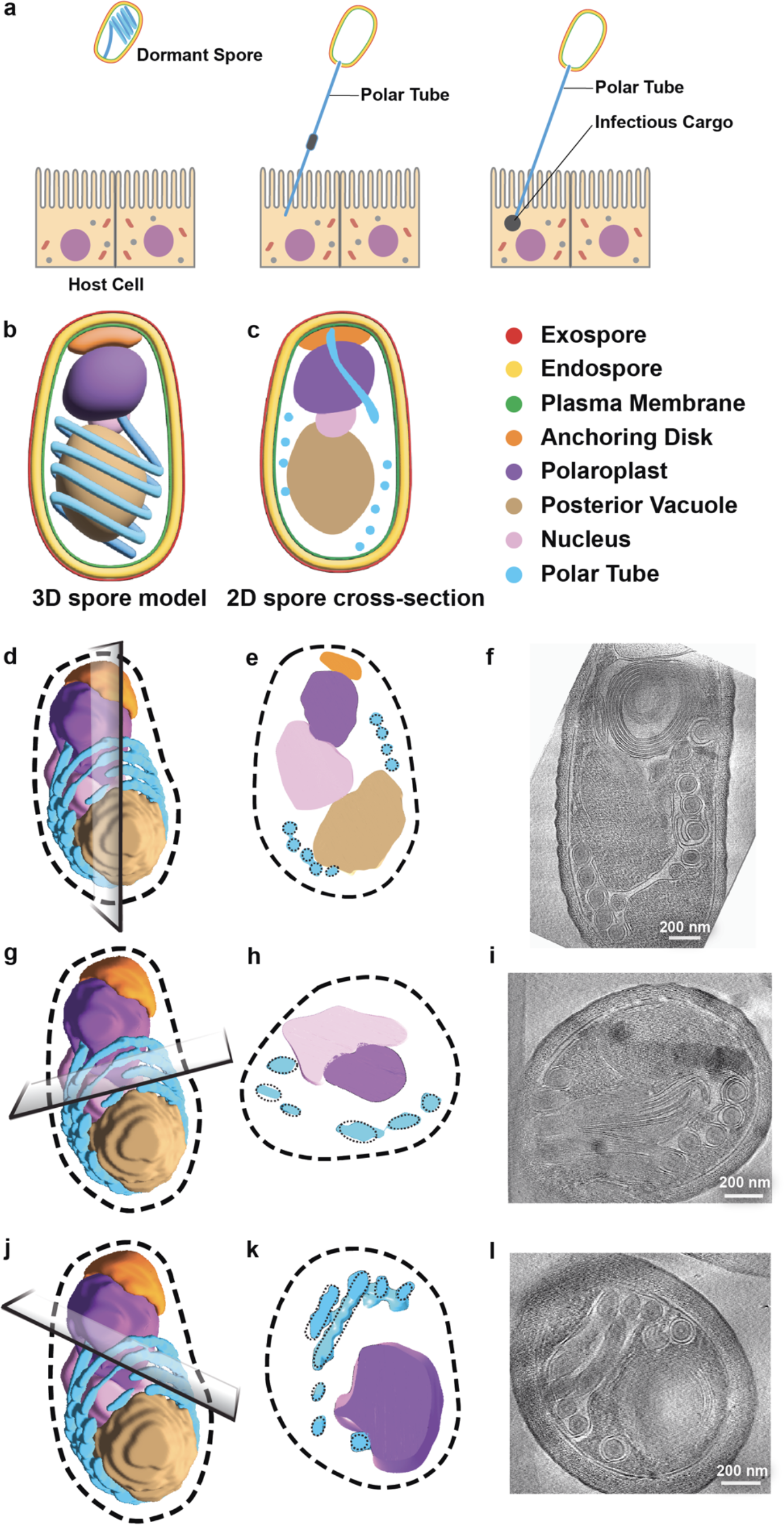
Correlation of SBF-SEM reconstructions and cryo-ET data to orient tomograms. **(a)** Schematic illustrating the microsporidian invasion process. The spore fires the long polar tube into the host cell, which transfers infectious cargo called the sporoplasm in the host. This process initiates infection. **(b)** Illustration of a microsporidian spore based on SBF-SEM data^51^. **(c)** Longitudinal cross-section of the illustration shown in (b). Color key applies to all parts of this Figure. SBF-SEM reconstructions are shown with the potential plane representing the lamella from tomograms (**d, g, j**), followed by a representation of the cross-section after slicing SBF-SEM data (**e, h, k**). 2D slices through the corresponding tomograms are shown in **f, i** and **l**. Correlating the cryo-ET data for each tomogram with cross-sections through the SBF-SEM model allowed us to orient tomograms and assign organelles.

In recent years cryo-electron tomography (cryo-ET) has emerged as a revolutionary technique that has allowed cell architectures to be studied in native, hydrated states in three dimensions (3D). Here, we employ cryo-ET to study the structural cell biology of the human-infecting microsporidian species, *Encephalitozoon intestinalis*. We focus on dormant spores, in which the PT is coiled in its pre-fired state, and the polaroplast is neatly organized at the anterior end of the spore. Our work sheds light on the architecture of the PT and associated organelles, including the polaroplast, in their cellular context. Combining cryo-ET with cellular modeling and previously published volume-EM data^51^, we propose integrative 3D models for the structure of these divergent organelles, and for the complex, interconnected membrane network inside the spore.

## Results

### Cryo-ET of dormant spores

To study the ultrastructure of dormant microsporidian spores, we vitrified purified spores, milled thin lamellae, and obtained cryo-ET data for the human-infecting microsporidian species, *E. intestinalis* (**Extended Data Fig. 1a**). In total, we collected 185 tilt series for *E. intestinalis* spores (**Supplementary Tables 1, 2**) and manually curated our dataset to remove low quality tilt series (48/185) and tilt series with lower contrast for internal spore features (49/185; see Methods; **Extended Data Fig. 1b-g**), resulting in a final dataset of 88 high-quality *E. intestinalis* tomograms. In addition, we analyzed 9 out of 12 tomograms from a previously collected dataset^52^ on dormant spores from the related species, *Encephalitozoon hellem* (**Extended Data Fig. 2, Supplementary Tables 1, 2**).

The tomograms are feature-rich and contain high-resolution snapshots of organelles within microsporidian spores. As each tomogram represents a ∼90-200 nm thick slice through the spore at a random angle, it can be challenging to relate the position and orientation of each tomogram to the whole spore, which is essential for assigning the identity of observed organelles and for developing a 3D model of a spore at higher resolution. To identify tomogram orientation, we visually correlated each tomogram with previously obtained volume-EM data^51^. We cropped a 3D reconstruction of an *E. intestinalis* spore (from Serial Block Face Scanning Electron microscopy; SBF-SEM; **Fig. 1b-i**)^27,51^ in a variety of different orientations to simulate the thin sections imaged by cryo-ET, and determined which orientation of the 3D reconstruction best recapitulated the orientation observed in each tomogram. Using the relative orientation within the spore, together with previous descriptions of organelle physical appearance from TEM^53–55^, we assigned the identity of most organelles in the spore.

### Overall organization of dormant spores

After correlating our tomograms to volume-EM data, we analyzed organelles from all available tomograms, and segmented organelles in 6 tomograms from *E. intestinalis* (**Fig. 2a, b, Extended Data Fig. 3, Supplementary Movies 1-5**) and 3 tomograms from *E. hellem*. This analysis sheds light on several aspects of the dormant spore, as outlined below.

**Figure 2.**
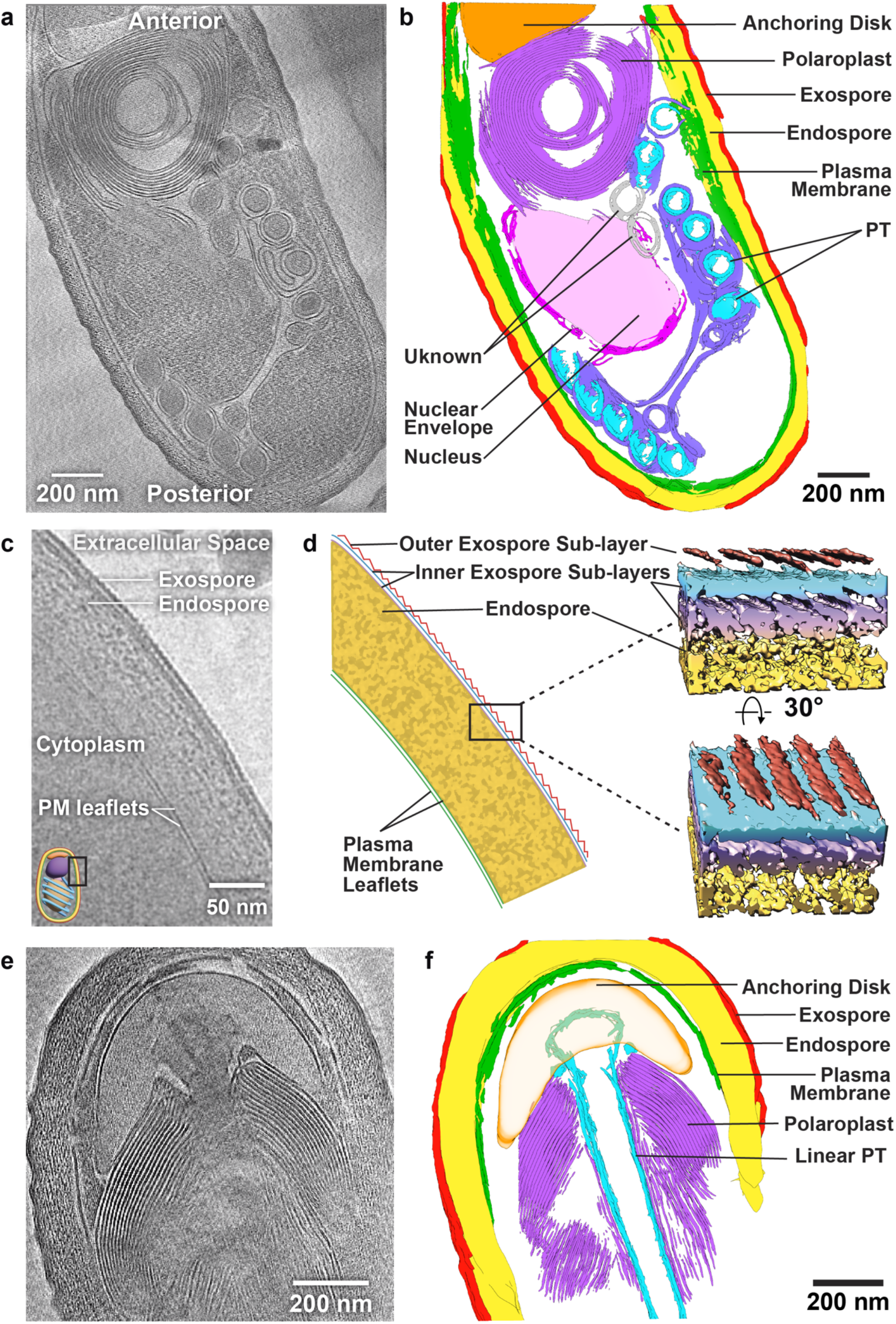
Overall organization of dormant microsporidian spores. **(a)** 2D tomogram slice through a representative spore. **(b)** 3D reconstruction of the spore shown in (a) after segmentation. **(c)** 2D tilt image showing the cell envelope of a spore, including plasma membrane, endospore and exospore, with saw-toothed pattern. **(d)** Schematic representation of tilt image shown in (c) and subtomogram average of the exospore (Gaussian filtered to 0.5Å and hide dust = 17). **(e)** Slice through a tomogram showing the linear part of the PT tucked in the anchoring disk and surrounded by polaroplast membranes. **(f)** 3D reconstruction of the spore shown in (e) after segmentation.

#### Cell envelope

The cell envelope of the spore comprises two major components: the plasma membrane and the spore coat (**Fig. 2c**). In some regions, the two leaflets of the plasma membrane are well resolved, with a bilayer thickness of 5 +/-0.2 nm. In almost all spores, we observe invaginations throughout the plasma membrane, as well as areas in which the density for the plasma membrane weakens (**Extended Data Fig. 4a**), which may represent connections of the plasma membrane to other organelles.

Similar to the fungal cell wall, the microsporidian spore coat comprises two distinct layers, the exospore and the endospore^56^ (**Fig. 2c, d**). The spore coat can either be smooth or have a ruffled appearance (**Extended Data Fig. 4b**), which does not clearly correlate with any other spore features, such as endospore thickness (**Extended Data Fig. 4c**). While the cause of spore coat ruffling remains unclear, factors such as turgor pressure or tension within a given spore may influence cell wall structure. The proteinaceous exospore is analogous to the outer mannoprotein layer in fungi^57^, and appears as an electron-dense outer layer ∼6-10 nm thick. Three sublayers of the exospore are observed, and in many spores a repetitive saw-toothed pattern is present on the outermost sublayer in some regions. 2D classification of single-tilt images, as well as subtomogram averaging (STA) from these spores, suggest that the outermost sublayer of the exospore may consist of a repetitive, protein-based array that forms a well-structured shell (**Fig. 2d, Extended Data Fig. 4d, e**). Sandwiched between the exospore and the plasma membrane is the endospore, which is analogous to the polysaccharide layer in fungi^58^, and is composed of glucans and chitin. Compared to the relatively thin exospore, the endospore is thicker (mean 68 nm +/-16 nm; range 40-115 nm), more irregular, and spongy in appearance (**Fig. 2c**).

#### Polar Tube, Anchoring Disk and Polaroplast

The PT is composed of a linear portion emanating from the anterior end of the spore, and a series of coils, found in the posterior half of the spore (**Fig. 2a, b, e, f**). The PT, observed in different orientations, is easily identifiable in most tomograms. In most cases, we observe the coiled region of the PT as circular or ellipsoidal cross sections (**Extended Data Fig. 5a**), but the linear portion of the PT (**Fig. 2e, f**) was also captured. At the anterior end, the linear portion of the PT flares into a funnel-like structure that is tucked inside the anchoring disk (**Fig. 2e, f**), an organelle that anchors the PT to the spore wall. Together, the linear portion of the PT and anchoring disc resemble a mushroom, with the PT forming the “stalk” and the anchoring disc forming the “cap”. The anchoring disk is bound by a single membrane, is electron-lucent, and is otherwise featureless. Across 8 tomograms, we observe a uniform gap of 34-37 nm between the anchoring disk and the plasma membrane, suggesting that specific protein complexes may span this gap.

Surrounding the linear portion of the PT is the polaroplast, which consists of numerous tightly stacked membranes^54^. We observe the polaroplast very clearly in at least 36 tomograms (**Fig. 2a, e**), and our cryo-ET data provide higher resolution information on the polaroplast than has been seen thus far. The membrane layers of the polaroplast are usually packed tightly against each other, with consistent luminal and cytoplasmic spacing (10.7 +/-1 nm), which may indicate the existence of protein complexes in the spaces in between. The packing of membranes in the polaroplast is visually reminiscent of the packing between membrane discs in retinal photoreceptor cone cells, where tetraspanins and other protein complexes help to organize and stabilize the membrane stacks^59^.

#### Nucleus, Vacuole and Mitosomes

The nucleus can be unambiguously identified in 16 tomograms, based on its size, double membrane structure, dense interior, and its mid-cellular location, where it nestles among the anterior-most coils of the PT (**Fig. 2a, b; Extended Data Fig. 5b**). The thickness of the nuclear envelope ranges from 7-20 nm, slightly thinner than in yeast (25-30 nm)^60^. Gaps are present in the nuclear envelope (**Extended Data Fig. 5b)** which may house nuclear pore complexes^60^. In 6 tomograms we can clearly observe double-membraned vesicular structures ranging from ∼ 57 nm to 102 nm in diameter, often proximal to the nucleus, and consistent with reported values for mitosomes (**Extended Data Fig. 5c**)^61^. Towards the posterior end of the spore, we identified two additional membranous structures: the posterior vacuole (**Extended Data Fig. 5d**) and a loosely packed membranous structure that may be an extension of the polaroplast **(Extended Data Fig. 5e**). The posterior vacuole is a large, electron-lucent, membrane-bound structure. Despite its large size and distinctive appearance, we do not always observe the vacuole. We suspect that while it is large and readily apparent in some spores, in others it may be much smaller.

### Structural insights into the polar tube

Our cryo-ET data allows us to define new architectural features of the PT. In our tomograms, PT cross-sections appear as concentric layers (**Extended Data Fig. 6a**), consistent with previous data from TEM sections of dormant spores from *Anncaliia algerae*^*49*^, *E. intestinalis*^*53*^ and *Enterocytozoon bieneusi*^*53*^. Two layers are particularly well contrasted, with density for additional internal layers observed in some coil-sections (**Fig. 3a, Extended Data Fig. 6a**). The outermost layer appears visually similar to the nearby polaroplast and plasma membranes. Therefore, we term this the Membrane (M)-layer. The M-layer surrounds a second layer of the PT. Unlike the relatively smooth and uniform appearance of the M-layer, the inner layer consists of regions with punctate, regularly-spaced, protein-like densities (**Fig. 3a**), and smoother, less punctate density around the remaining circumference of the layer. To better understand how the punctate densities are organized in 3D, we performed segmentation analysis, in which we marked the center of mass of each observed density on each slice of a tomogram (**Fig. 3a**). When unable to decipher puncta, we segmented the circumference of this layer as a single object. The resulting 3D reconstructions clearly show that the punctate densities correspond to a series of parallel filaments that spiral around the PT in a left-handed helical arrangement (**Fig. 3b, Extended Data 6b**). Based on the presence of protein-like filaments, we name this the Outer Filament (OF)-layer. In some areas of the OF-layer, filaments could not clearly be resolved, which may be explained by the orientation of a given PT cross-section relative to the plane of the lamella and to the tilt axis of the microscope (see Methods; **Extended Data Fig. 6c**). Thus, we propose that filaments are likely distributed around the entire OF-layer. The filaments are regularly spaced, approximately 65.5 +/-5.2 Å(n=9) apart, and are tilted -25.3° +/-3.8° (n=9) on average relative to the long axis of the PT (**Extended Data Fig. 6b**). To estimate filament numbers within the OF-layer, we focused on 7 PT coil sections within a single tomogram (tomogram 4), and measured the circumference of the OF-layer for each section, which varied from 284 nm to 319 nm. Based on the circumference of each coil, and the inter-filament spacing and tilt angle, we calculated the expected number of filaments per coil section, which ranges from 37 to 44. Interestingly, the range in filament number suggests the possibility of a variable number of filaments along a single tube. A range in filament number along the polymer is not entirely unprecedented, and is commonly observed in microtubules, in which different protofilament numbers can be observed even within the same microtubule^62,63^. The OF-layer forms a continuous protein shell, which may serve as a skeleton that gives the PT its overall shape (**Fig. 3b**).

**Figure 3.**
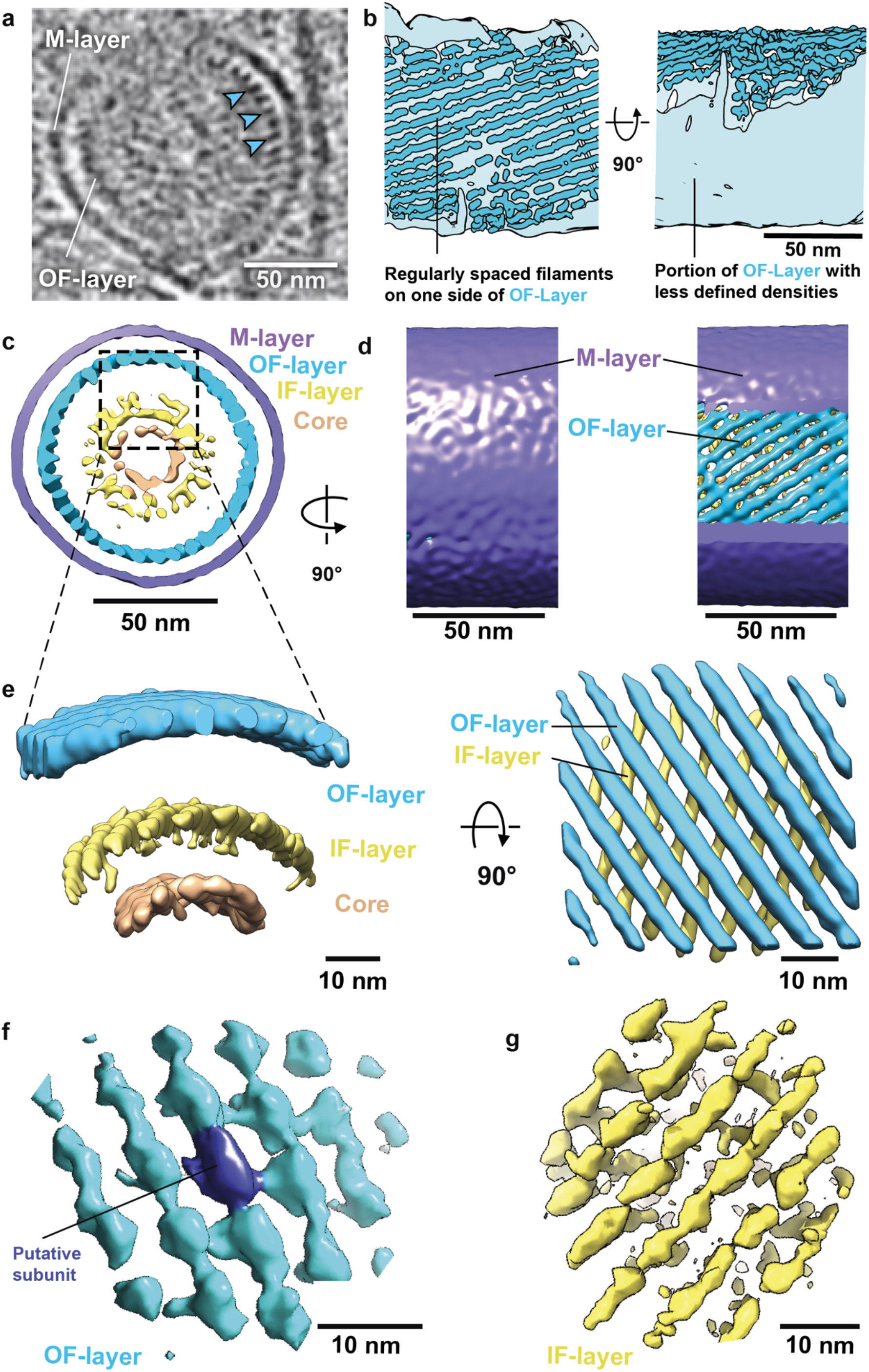
Architecture of the polar tube from dormant spores. **(a)** A single PT coil cross section from a representative tomogram. Layers of the tube are clearly visible. Cyan arrowheads indicate consecutive discrete densities observed in the OF-layer of the tube. **(b)** 3D reconstruction of the OF-layer after segmentation, in which discrete densities appear as filaments along the portion of the tube. Two orientations are shown, one in which an arc of segmented, discrete densities appear as filaments, and the other regions of the tube, in which discrete densities were not discernible, and appear as a smooth layer. As discussed in the text, this is likely due to technical limitations. Subtomogram averaging was performed on the whole PT coil sections, shown as a cross-sectional view **(c)** and side view **(d)**. The side view in **(e)** is shown with the M-layer peeled back to reveal left-handed filaments in the OF-layer (right). **(f)** Focused subtomogram averaging of different layers in the PT. Focused refinement of the OF-layer **(f)** and the IF-layer **(g)** show the presence of regularly spaced repeating units in the OF-layer and clear filaments with hints of a repeating unit in the IF-layer.

Most PT coil-sections were individually wrapped by the M-layer. However, in 11 out of 82 *E. intestinalis* spores, 2 or more OF-layer cross-sections were encapsulated within a shared M-layer (**Extended Data Fig. 6d**). Some tomograms included OF-layer cross-sections surrounded by a shared M-layer alongside individually wrapped OF-layer cross-sections (**Extended Data Fig. 2a, Extended Data Fig. 6d**). One possible model consistent with these observations is that, earlier in spore development, the entire PT assembles within a membrane compartment (**Extended Data Fig. 6d**). As the spore matures, membrane remodeling leads to the compartmentalization of each PT coil, resulting in OF-layer cross-sections that are pinched off and individually wrapped by the M-layer.

To gain further insights into the structure of the PT, we used subtomogram averaging, which has the advantage of boosting signal to noise to generate 3D reconstructions at higher resolutions (**Supplementary Table 3**). We selected ten spores based on data quality, high signal-to-noise, and number of PT cross-sections in the field of view. We performed subtomogram averaging of the whole tube, in which all sections of the tube from a given spore were averaged together (**Extended Data Fig. 7, 8**). Attempts to average across different spores were unsuccessful, possibly due to structural heterogeneity between spores. Consistent with segmentation analysis, subtomogram averaging reveals a smooth, featureless M-layer with a diameter of 114 +/-7.7 nm. The M-layer surrounds the OF-layer, which has a diameter of 89 +/-4.6 nm, and is formed by left-handed filaments spaced ∼60 Å apart (**Fig. 3c-e, and Extended Data Fig. 8a, b**). In the subtomogram averages of the PT from six of the 10 spores analyzed, the resolution was sufficient to estimate the number of filaments in the OF-layer by counting, albeit with modest uncertainty in some regions (**Extended Data Fig. 8a**). Consistent with the segmentation analysis from a single tomogram, we count 36-41 filaments around the OF-layer (**Supplementary Table 3**).

We performed focused refinements on the OF-layer, which improved the definition of filaments, and allowed us to analyze these structures with more confidence (**Fig. 3e, Extended Data Fig. 8c, d, Supplementary Table 4**). In our best map, we observe the presence of a possible structural unit repeating along the filaments with a periodicity of ∼58 Å (**Fig. 3f**). The filaments may be composed of one or more proteins such as PTP1, PTP2, PTP3 and/or PTP6, which have previously been identified as being localized along the length of the PT^6,42,43^. Based on the apparent volume of the repeating unit, we estimate that it would correspond to a globular protein with a molecular weight of ∼50-90 kDa. This could represent homo-or hetero-oligomers of PTP1, PTP2, and/or PTP6; however, PTP3 is substantially larger at ∼137 kDa.

In addition to the M- and OF-layers, subtomogram averaging of the whole tube revealed additional density at the center. On careful inspection, this density can sometimes be observed in the raw tomograms as well (**Extended Data Fig. 6a**). Local refinement of this central density further resolved two parts: 1) an additional discrete layer consisting of filaments, which we call the Inner Filament layer (IF-layer); and 2) a central Core (C)-region. The IF-layer is ∼51 +/-3.2 nm in diameter and is formed from filaments with a right-handed twist, opposite to the left-handed filaments of the OF-layer (**Fig. 3e, g, Extended Data Fig. 8d, e, Supplementary Table 5**). The inter-filament spacing in the IF-layer is similar to what was observed for the OF-layer, ∼60 Å (**Fig. 3g and Extended Data Fig. 8d, e**) and the angle of the filaments relative to the longitudinal axis of the PT is ∼25°. Assuming consistent spacing, we estimate there are ∼24 filaments in the IF-layer. At the center of the tube, the C-region, which is ∼39 +/-4 nm in diameter, shows clear density for possible additional proteins, but lacks clearly defined features, and is separated by a ∼8 nm gap from the IF-layer.

Together, our structural analysis of the PT revealed four distinct components: M-layer, composed largely of membrane; OF-layer, composed of left-handed filaments; IF-layer, composed of right-handed filaments; and the C-region, at the core of the tube.

### 3D topology of the polaroplast

The PT is closely associated with the membranous polaroplast organelle, unique to microsporidia^30^. Our cryo-ET dataset includes 36 tomograms in which the polaroplast is visible in different orientations, affording us the opportunity to study this organelle in 3D. The complex nature of the polaroplast was apparent when comparing its appearance among different tomograms. In some cases, it appears as a series of concentric rings of membrane (**Extended Data Fig. 9**), resembling the layers observed when slicing through an onion. In other cases, the polaroplast appears to consist of closely-spaced sheets of membrane (**Extended Data Fig. 9**), forming large stacks reminiscent of light sensing organelles, such as the flattened membrane disks in the outer segment of photoreceptor cells^59^, or in thylakoid membranes^64^. In some cases, more complex folds and whorls are apparent (**Extended Data Fig. 9**). Given that each tomogram represents only a relatively thin slice through the spore at an arbitrary angle, and that the position and structure of membranous organelles may vary from one spore to the next, conceptualizing the 3D organization of the polaroplast based upon the limited depth of each tomogram is challenging. In addition, the 3D reconstructions of entire spores from volume-EM data^51,27^ are too low in resolution to resolve these intricate membranes and inform on this question. We therefore turned to 3D cellular modeling to generate and refine hypotheses for how the polaroplast may be organized. We generated six virtual models for the 3D organization of the polaroplast (**Fig. 4a**). These models differ from one another in three main ways: 1) whether the PT is completely encircled by the polaroplast or not; 2) whether stacks are formed from membrane folding back-and-forth, or from membrane spirals; and 3) whether the axis of membrane folding/spiraling is parallel to the long axis of the PT straight segment, or is orthogonal (**Fig. 4a and Supplementary Table 6**). By cropping our six candidate 3D models at various angles, and correlating their appearance with the appearance of the polaroplast in a subset of tomograms, we found that some models were better able to account for the experimental data than others (**Fig. 4a, b, c, Extended Data Fig. 10**) (see Methods, **Supplementary Table 7 and Supplementary Table 8**). The two models that best account for the experimental data either fold back and forth (model 5) or spiral (model 2) the membrane about an axis parallel to the PT to yield a stack of membrane sheets, and the stack curves to cradle the PT but only partially encircles it. We judged model 5 to be a somewhat better fit than model 2, although the differences were relatively minor and we consider both models to be in fairly good agreement with experimental data. Differentiating between Model 5 (membrane folding) vs. Model 2 (membrane spiraling) would require tomograms that catch very specific views of the polaroplast with sufficient clarity to trace membrane connectivity unambiguously between the layers. In several of our tomograms, key regions of the membrane topology in the tomogram were difficult to interpret, leaving some ambiguity in the model (**Fig. 4, Extended Data Fig. 10**). What became apparent from the 3D cellular modeling is that if the model is sliced and rotated in different orientations, the appearance of the polaroplast is strikingly different. Thus, the close interplay between 3D modeling and experimental data allowed us to elevate our understanding of the cryo-ET data and propose a model for the 3D organization of the polaroplast organelle in dormant spores.

**Figure 4.**
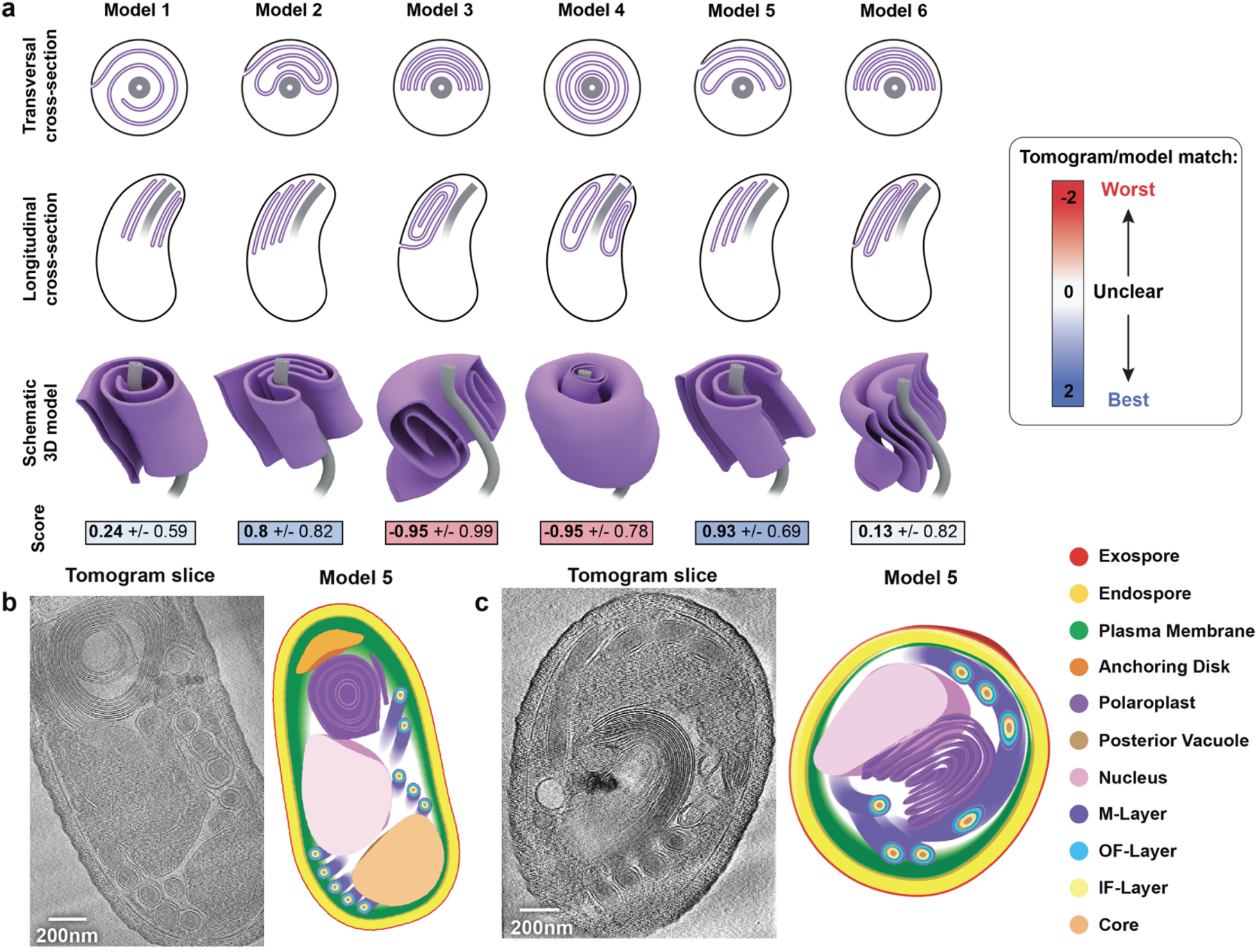
Architecture of the polaroplast in dormant spores. **(a)** Schematic showing six virtual models generated to study possible polaroplast organization. Scores indicate the visual agreement between each model and a subset of tomograms, scored as described in Methods. **(b, c)** Examples of two tomograms with the corresponding view of the best-matching model (model 5) from the six models shown in (a). (b) uses the short polaroplast model whereas (c) uses the long polaroplast model.

The origin of the polaroplast and its relation to other membranous organelles within the spore remain unclear. Intriguingly, in two tomograms, we observed connections between the polaroplast and other cellular membranes. First, in a tomogram of an *E. hellem* spore, the polaroplast membrane is clearly continuous with the PT M-layer, suggesting that the M-layer is likely derived from the polaroplast membrane network. In this tomogram, the polaroplast connects to an M-layer encapsulating four PT cross-sections (**Fig. 5a, Supplementary Movie 6**), alongside a single individually wrapped PT segment. Second, one of our *E. intestinalis* tomograms reveals a physical connection between the polaroplast and the plasma membrane: an invaginated pocket of plasma membrane is continuous with the polaroplast membrane stack (**Fig. 5b, Supplementary Movie 7**). These physical connections suggest that the plasma membrane, polaroplast, and PT M-layer together may form a single, highly complex membrane system, and that the lumen of the polaroplast and PT structures within the M-layer may be topologically continuous with the extracellular environment. This polaroplast– plasma membrane–M-layer network model supports the idea that the proteins that make up the PT reside within an extra-cytoplasmic compartment: the structural components of the PT may be topologically outside the plasma membrane but inside the spore coat (**Fig. 6a**), consistent with the predicted N-terminal secretion signals on PTPs^45,65,66^ and previous hypotheses^49^.

**Figure 5.**
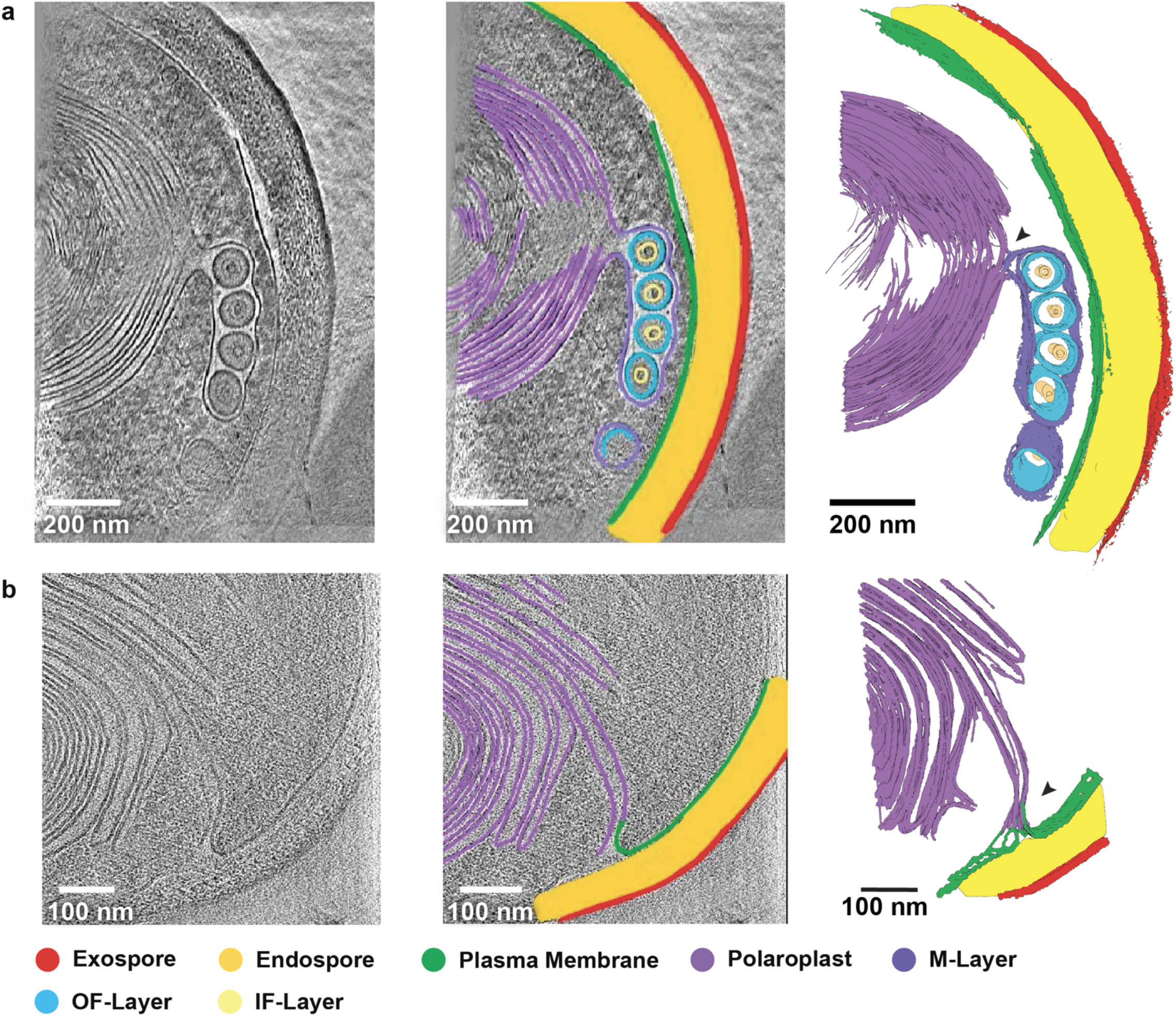
Membrane connectivity in dormant spores. **(a)** Tomogram slice, annotated tomogram slice and 3D reconstruction from segmentation, showing the connection between the polaroplast and the M-layer of the PT (black arrowhead). **(b)** Tomogram slice, annotated tomogram slice and 3D reconstruction from segmentation, showing the connection between the polaroplast and plasma membrane (black arrowhead).

**Figure 6.**
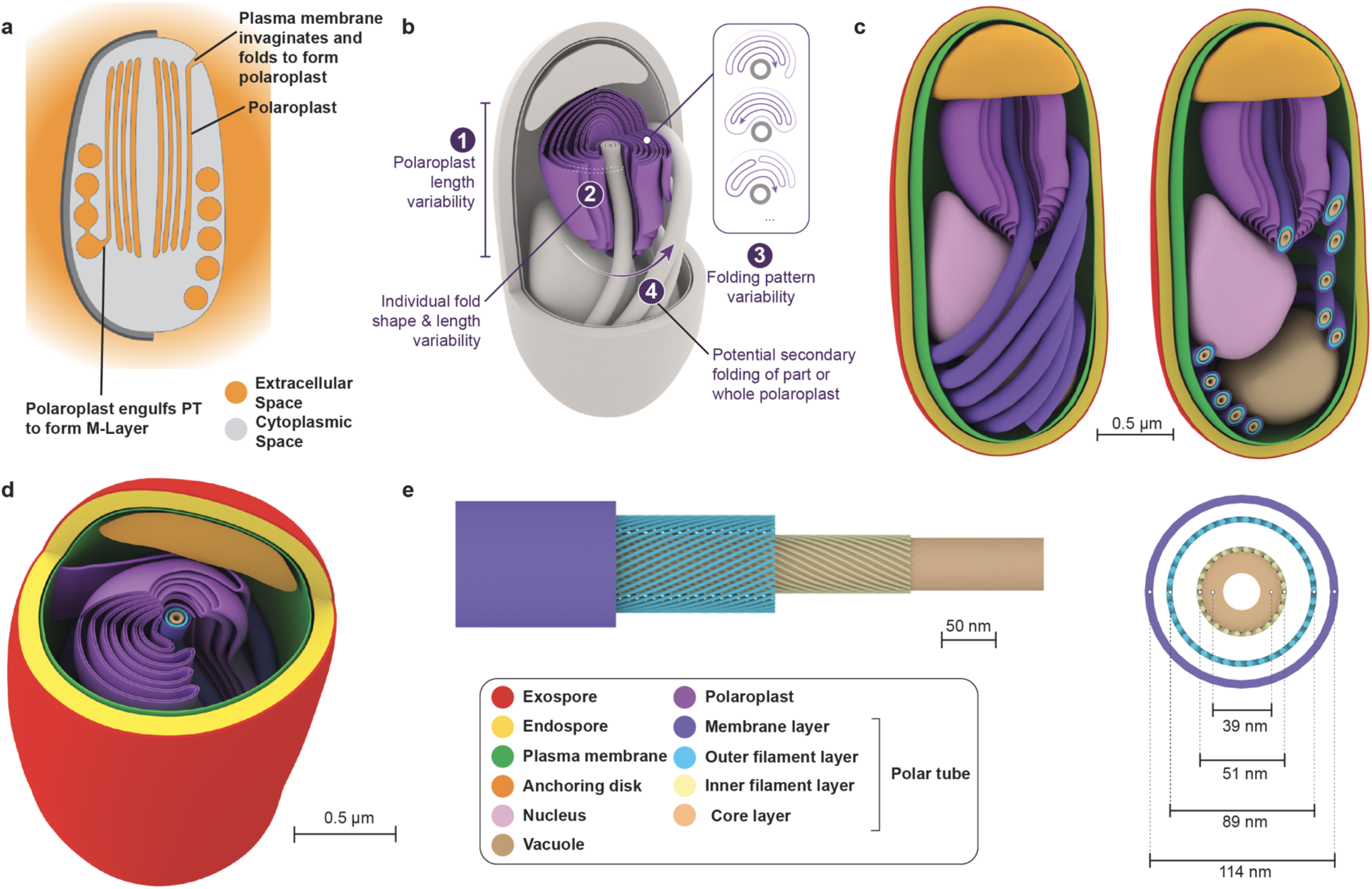
Structural model of a dormant microsporidian spore. **(a)** Schematic highlighting topology of the microsporidian PT. Orange color indicates topologically extracellular material. Based on the connection between the polaroplast and the plasma membrane, and the connection between the polaroplast and M-layer of the PT, the PT and polaroplast are shown as being topologically extracellular. **(b)** Model for the organization of the polaroplast. Model 5 was chosen as it agrees best with the cryo-ET data. Variations of the polaroplast organization that was observed in cryo-ET data are highlighted in this model. This model represents our current understanding of the polaroplast architecture in 3D, together with variations that were observed. **(c, d)** Model of the spore highlighting organization of the various organelles, shown as side (c) and top (d) views. **(e)** Model for PT structure, highlighting the M-layer, OF-layer, IF-layer and core.

## Discussion

### Integrative modeling to study spore organelle architecture

The unique cellular organelles of microsporidian spores, coupled with the limited observable volume in the cryo-ET lamellae, made assigning organelles and extrapolating the relative orientation of the tomogram challenging. To overcome this, we manually correlated volume-EM data with cryo-ET data for *E. intestinalis* spores, which allowed us to confidently assess the cutting plane within the cell during lamella preparation. While this process was very helpful, it was laborious, and necessarily involved a certain amount of subjectivity in correlating the data. Automated correlation of volume-EM and cryo-ET data with clearly defined metrics for assessment, such as cross correlation, may be helpful to incorporate in structural cell biology workflows. One caveat is that this kind of correlation is only applicable for cells in which organelles are oriented in a consistent spatial arrangement relative to other organelles.

The polaroplast is a highly divergent membrane-rich organelle only described in microsporidian species, the 3D organization of which was not well understood. While our cryo-ET data provided excellent resolution for tracing the membranes and interactions with nearby organelles, the limited volume of our lamellae initially prevented us from conceptualizing how the whole organelle is structured and packed inside the spore. Piecing individual tomograms together to generate a 3D model at the structural cell biology level required us to incorporate cellular modeling. Based on initial observations in the cryo-ET dataset, we determined the three criteria described above **(Fig. 6b)**, leading to six possible topological models for the polaroplast. To introduce rigor, one author prepared guidelines for scoring that were shared with five authors, who blindly scored each model against tomograms (**Extended Data Fig. 10**). This process resulted in five sets of independent and blind scores assessing the models, which the authors discussed and which were generally consistent. Cross-sections of model 5, showing horizontal folding matched well with almost all tomograms (**Extended Data Fig. 10**), and thus is currently our favored model for the structure of this organelle (**Fig. 6c, d**). While it is now becoming feasible, and will soon be standard to collect large cryo-ET datasets, the interpretation of the cryo-ET data to answer biological questions requires integration of other approaches. In this case, cellular modeling was key to going from a description of tomograms to a 3D model. Certainly, this model will be refined in the future as more data become available. The structure of the polaroplast, coupled with our finding that it is continuous with the M-layer of the PT provide a physical basis for the long-standing hypothesis that the polaroplast is the source of membrane that encases the infectious cargo being transported through the PT^35^.

### Structural and mechanistic insights into the PT

Integrating our segmentation and subtomogram averaging analysis of cryo-ET data, we conclude that the PT in the dormant spore consists of at least four main layers: a membranous M-layer which surrounds the whole tube, an OF-layer, followed by an IF-layer, both composed of regularly spaced filaments with opposite handedness, and a central core structure that currently is only partially resolved (**Fig. 6e, Supplementary Movie 8**). We previously reported that the PT always forms a right-handed coil within the spores of *E. intestinalis* and two other microsporidian species^27,51^. Interestingly, the asymmetric, arrangement of the filaments that make up the OF- and IF-layers provide a possible explanation for the right-hand bias of the PT coil in spores: just as the right-handed molecular organization of DNA leads to preferential formation of left-handed supercoils^67^, so the asymmetric arrangements of the PT building blocks may dictate the microscale organization of the tube in the spore by favoring right-handed coiling. While it is fairly common for larger scale coiling to take the opposite hand of the twist of the building blocks, with molecules such as DNA as well as macroscale objects such as rope, opposing handedness of the OF- and IF-layers complicate this simple interpretation. Further studies will be needed to understand the relative contributions of the OF- and IF-layers to the mechanical properties of the PT, and its larger-scale organization.

Based on previous lower resolution studies of the post-firing PT^6,44,48,68^, the protein component of the PT was described to form a hollow tube, well-suited for transporting cargo. Intriguingly, a recent cryo-ET study^44^ of the post-firing state of the PT from another microsporidian species, *V. necatrix*, suggested that the outermost layer may be protein, with a membranous inner layer, the opposite of what we observe in the pre-firing state. It is likely that during the firing process, major reconfigurations of the PT occur to facilitate cargo transport. In our pre-firing PT reconstructions, the presence of the IF-layer and core within the OF-layer is somewhat confounding: the function of the PT is to transport cargo, yet inside the spore the lumen of the PT is blocked. It is likely that at least the core, and possibly the IF-layer, exit the lumen within the OF-layer during firing. While the mechanism of PT firing is still uncertain, it is generally believed that at least part of the PT everts or turns inside out while firing^28,69–71^. However, these ideas have not accounted for the complexity of PT layers and structure. One possibility is that the core and/or IF-layer are ejected during the process, and may be involved in piercing the host cell, delivering protein toxins or modulators to the target cell, or may be released into the surrounding media. After the ejection of the core and/or IF-layer, the OF-layer may evert, resulting in a hollow tube for transport. Future work focused on high-resolution and time-resolved analysis of PT firing from the same species will shed light on the mechanism of PT firing.

## Acknowledgements

We thank Elizabeth Villa, Joshua Hutchings, Tom Laughlin, members of the Villa Lab, Garrett Greenan, Zhen Chen, Reza Paran, Jake Johnston for helpful discussions and advice in initial cryo-ET data processing. We thank Noelle Antao, Fred Rubino, Alice Herneisen, Paris Watson and Kacie McCarty for their critical reading of and valuable feedback on this manuscript and all members of the Bhabha+Ekiert Labs for helpful discussions. We gratefully acknowledge the following funding sources. Award number 915749 (American Heart Association to M.U.), SSP-2018-2737 (Searle Scholars Program, to G.B.), R01AI147131 (NIAID, to G.B.), Irma T. Hirschl Career Scientist Award (to G.B.). G.B. is a Pew Scholar in the Biomedical Sciences, supported by The Pew Charitable Trusts (PEW-00033055). Some of this work was performed at the National Center for In-Situ Tomographic Ultramicroscopy (NCITU) and the Simons Electron Microscopy Center located at the New York Structural Biology Center, supported by the NIH Common Fund Transformative High Resolution Cryo-Electron Microscopy program (U24 GM129539,) and by grants from the Simons Foundation (SF349247) and NY State Assembly. Some of this work was performed at NRAMM. The National Resource for Automated Molecular Microscopy is supported by a grant from the National Institute of General Medical Sciences (9 P41 GM103310) from the National Institutes of Health. Cryo-ET data processing was performed using computing resources at the HPC facility at NYU, and at the Advanced Research Computing at Hopkins (ARCH) core facility (supported by the National Science Foundation grant number OAC 1920103). and we thank the members of the HPC teams for computing support.

## Data Availability

The cryo-ET data, with segmentations were deposited in the EMPIAR (Electron Microscopy Public Image Archive) database under the accession codes: EMPIAR-12176 (Dataset 1 and 2), EMPIAR-12177 (Dataset 3), and EMPIAR-12175 (Dataset 4). The cryo-ET data, with segmentations were deposited in Cryo-ET Data Portal built by the Chan Zuckerberg Imaging Institute and Chan Zuckerberg Initiative with deposition ID CZCDP-10307, which contains Dataset 10436 (corresponding to 1 and 2), Dataset 10437 (corresponding to Dataset 3), and Dataset 10438 (corresponding to Dataset 4). The polar tube, polaroplast and spore models described in this manuscript are also deposited in Github at (https://github.com/mahrukhu/Polaroplast-Models).

The best subtomogram averages have been deposited in the EMDB (Electron Microscopy Data Bank) database under accession codes: EMD-45674 (whole polar tube), EMD-45671 (OF-layer), EMD-45672 (IF-layer), and EMD-45673 (OF and IF layers).

## Author contributions

M.U., G.B. and D.C.E. conceptualized the project. M.U., D.B., and M.K. collected data. M.U., N.C., M.R., R.R., H.R., E.D.Z., J.H.I., D.C.E. and G.B. performed data analysis. M.U., N.C., D.C.E. and G.B. wrote the manuscript. All authors provided feedback and edited the manuscript.

## Methods

### Spore propagation and purification

*E. intestinalis* microsporidian spores (ATCC: 50506) and Vero cells were maintained in Dulbecco’s Modified Eagle Medium (DMEM) (Gibco 11995065), supplemented with 10% heat-inactivated fetal bovine serum (FBS) (VWR 89510-188) and 1X MEM-NEAA (Non-essential Amino Acids) (Gibco 11140050), and were grown at 37°C, 5% CO_2_. At 90% confluency the media was replaced with DMEM supplemented with 3% FBS and 1X MEM-NEAA and a single vial of spores were added. Vero cell maintenance and spore propagation were carried out in 75 cm^2^ tissue culture flasks and the media was replaced with fresh media every two days.

Spores were harvested 14 days post-infection. Cells were detached from flasks using a cell scraper and the suspension was centrifuged at 1800 *x g* for 10 minutes at room temperature. Supernatant was discarded. Cells were resuspended in 1X DPBS (Gibco 14190144) and mechanically disrupted using a G27 needle to release any spores present inside cells. To purify spores, an equal volume of 100% percoll (Cytiva 17-0891-02) was added to the cell slurry in PBS and vortexed to homogenize the mixture. The solution was then centrifuged at 1,800 *x g* for 30 minutes at room temperature. The purified spore pellets were washed 3 times with 1X PBS and stored at 4°C until further use. Spores for dataset 3 were purified until this step and used for cryo-ET sample preparation. Dataset 4 referred to in this manuscript comes from previously collected and published *E. hellem* data^52^. For datasets 1 and 2, an additional purification step was carried out. The purified spore pellets were added to a discontinuous percoll gradient consisting of 25%, 50% 75% and 100% percoll, and centrifuged at 12,700 x *g* with a TH-641 (Thermo Scientific Cat # 54295) rotor at 25 °C for 30 minutes. The pellet fraction was collected and washed in 1X DPBS (Gibco 14190144) three times and stored at 4°C until further use.

### Cryo-ET grid preparation

The waffle method^52^ was used for grid preparation. Spore concentrations frozen per grid were as follows: 5.7 × 10^7^ spores/µL for dataset 1; 8 × 10^5^ spores/µL for dataset 2; 1.3 × 10^8^ spores/µL for dataset 3. 5 µL of sample was applied to each waffle grid assembly, and the sample was frozen using Wholwend Compact 01 high pressure freezer, with a pressure of 2,100 bar. All waffled grids were carefully clipped prior to milling^52^.

### cryo-FIB milling and lamella preparation for cryo-ET

Lamellae of 90 to 200 nm size for cryo-ET data collection were milled. Below is a detailed description of how milling was performed on grids for datasets 1, 2 and 3. Dataset 4, analyzed in this manuscript, was previously published^52,72^. We followed a published protocol for waffle grid preparation and milling^72^. All parameters were kept the same as in the published protocol unless otherwise stated. Milling was performed on Thermo Fisher Scientific Aquilos 2 cryo-FIB/SEM instrument at NCITU.

#### Preparation and lamella site assignment

For each grid that was milled, a new project was created in MAPS (Thermo Fisher Scientific) and an SEM grid overview was recorded. This was followed by platinum sputter coating for 15 seconds using a current of 30.0 mA and pressure 0.10 mbar. After the system was recovered from sputtering a layer of organometallic platinum (also known as GIS platinum) was added. With our milling settings, we used 2 minutes of GIS for lamellae being milled with 200 nm thickness and 2 minutes 15 seconds for lamellae being milled with 150 nm thickness. Using MAPS software, ROIs and preliminary lamellae sites were defined on the grid. The grid was then moved from the mapping position to be orthogonal to the ion beam (IB) which was kept at 30 KV throughout our protocol. The microscope was operated using xT software (Thermofisher Scientific).

#### Pre-cuts

With an ion beam current of 10 pA, we navigated to one of the milling sites marked in MAPS. An image was acquired and the contrast was adjusted such that the lamella site was clearly visible. The surface of each lamella site was inspected to ensure it was flat and free of contamination. To locate grid bars, small windows were milled using a current of 15 nA close to where they are expected to appear. Once located, we performed precuts away from the grid bars. Pre-cuts were performed using a ‘pre-cut’ pattern on the xT software with a current of 15 nA. Contrast was adjusted during this process to ensure that sufficient sample was removed with the beam. Before moving to the next lamella site, the current was changed back to 10 pA and the same steps were repeated on all lamella sites.

#### Pre-cleans

After all the precuts were completed, we returned to the mapping position and acquired a new tileset with the same parameters as before. Old preliminary lamella sites were deleted and new sites were assigned on the grid overview using the precut positions. The project was then opened in AutoTEM (Thermo Fisher Scientific) and a milling template (parameters as described in Klykov, Bobe et. al, 2022)^72^ was applied to all the lamellae sites. We then ran “preparation” for all lamella sites in guided mode and followed the prompts for eucentric tilt and a 20° milling angle. The lamellae were positioned to guide notch placement. This was followed by the pre-cleaning step which is performed in a stepwise manner. A milling angle of 45° is set with milling current 7 nA, followed by 30° with milling current 7 nA and finally 20° with milling current 3 nA.

#### Notch milling and final lamellae positioning

The notch pattern was placed at the edge of the slab of the pre-defined lamella site. The dimensions of the notch pattern as described in (Klykov, Bobe et. al, 2022)^72^. The notch was milled using a current of 0.3 nA. Final lamella placement was done in AutoTEM, where the lamella was placed 1 µm away from the notch. Automated milling and thinning of lamella sites was done in stepwise mode overnight using AutoTEM.

### Cryo-ET data collection and pre-processing

Tilt series were collected in 3 sessions on 3 different Titan Krios microscopes equipped with K3 cameras (**Supplementary Table 1**). We aligned and reconstructed tomograms from 185 tilt series from *E intestinalis* (dataset 1, 2 and 3 from **Supplementary Table 1**) and 12 tilt series from previously collected *E hellem*^*52*^ (dataset 4 from **Supplementary Table 1**). Data were collected at magnifications ranging from 26,000 to 42,000, corresponding to pixel sizes of 2.077 - 3.431 Å/pixel (**Supplementary Table 1**). The acquisitions were controlled either with Leginon^73^ or SerialEM^74^ with or without PACE-tomo^75^ (as noted in **Supplementary Table 1**), using 2 or 3 degrees step increments, a dose symmetric scheme and defocus ranges of -4 to -7 µm. At each tilt, movies with 10 to 23 frames were acquired, using dose per tilt ranging from 1.13 to 2.87 e^-^/Å^2^.

Datasets 1 and 4 were pre-processed using MotionCor2^76^ and protomo^77^, followed by tilt alignment and reconstruction with either IMOD^78^ or AreTomo^79^. Datasets 2 and 3 were motion-corrected in WARP^80^ and tilts were aligned with AreTomo^79^. For visual analyses, tomograms and tilt series were imported into FIJI^81^ or IMOD^78^. Images from reconstructed tomograms or tilt series shown in this manuscript are also exported from these programs.

To assess whether cell-to-cell variability with respect to contrast of internal organelle structures could be the result of a technical issues, we assessed the following for each tilt series: 1) ice thickness, as estimated using the dose rate per tilt extracted from SerialEM^74^ mdoc files and 2) defocus, estimated by EMAN2^82^. We found no correlation between internal cellular contrast and either of these two parameters. Importantly, we collected a single tilt series which contained both a spore with high contrast of internal organelles and a spore with low contrast of internal organelles within the same field of view (**Extended Data Fig. 1f**). These observations led us to believe that the differences we note represent biological heterogeneity, and are not the result of a technical issue.

### Segmentation

Tomograms with ID 15, 25, 46 and 90 (**Supplementary Table 2**) were segmented using an auto-segmentation method. Tomograms with ID 1, 3 and 104 were segmented manually (**Fig. 2b, f, 4d, e, Extended Data Fig. 3**). We used 8x binned, SART reconstructed tomograms as input for segmentation of overall spore structures such as PT, spore envelope, nucleus. To segment polaroplast membranes in tomograms 15, 90, 46 and 25 we used 2x binned tomograms. For tomograms that were auto-segmented, segmentation was performed using Dragonfly software, Version 2022.1.0.1259 for [Windows]. Tomograms were first preprocessed with a 3D Gaussian filter, histogram equalization, and an unsharpening filter. For a given tomogram, between 3 and 5 slices along the z-axis were manually annotated voxelwise for organelles of interest. The annotated slices were used to train a 2D U-Net model with DICE loss within the Segmentation Wizard tool. The trained model was then used to predict the segmentation for all z-slices in the tomogram. In some cases, a model trained on one tomogram was applied to a different tomogram, after which a few of the automatically segmented slices were manually corrected and used for re-training. For some tomograms, an additional model was trained in the Deep Learning Tool using a cross-entropy loss, with the weights increased for organelle classes that are rare or difficult to segment. Following segmentation of a full tomogram, noise was removed interactively, as well as by removing segmented components below a size threshold using the process islands function in Dragonfly. Finally, the predicted segmentation was overlaid with raw tomogram slices and any areas that were left unsegmented in the model were manually segmented. Manual segmentation on tomograms 1, 3 and 104 was also performed on Dragonfly where each organelle was identified and traced by the user in separate regions of interest. All segmented objects were converted into contour meshes and smoothed and visualized in chimeraX^83^. Discrete densities present on the OF-layer were segmented by placing a marker at the center of mass of each distinct density throughout the volume where that density was visible Discrete densities in the OF-layer can also be observed on raw tilt series where features can be more clearly seen on one side of the coil or the other depending on the tilt and on the orientation of the tube relative to the tilt axis and projection. It is possible that the filaments are present across the entire circumference of the PT coil sections but because of the way the spore is sliced, the densities representing the filaments are more clear in certain directions than in others. To test this, we generated a 3D model of a PT with filaments all the way around, and sliced and tilted it to mimic a tomogram. Depending on the orientation of the cross-section, one side of the PT (where the axis of projection aligns best with the orientation of the filaments) shows clear discrete densities, while the other side does not, despite the model containing filaments all the way around. As this observation is consistent with what we observe in our data, we conclude that filaments are likely all around the OF-layer.

### Correlation of SBF-SEM data with spore tomograms

A 3D reconstruction of a spore from EMPIAR-11683 was opened in Chimera^84^ and the shape of the spore section on the tomogram was used as a guide to orient the SBF-SEM in the longitudinal axis or transversal axis. This was then clipped from the front and back to produce a thin slice which was finely rotated to reproduce the orientation of organelles to match each tomogram.

### 2D classification of the cell envelope

Tilt series images were imported into Relion 4.0^85^ and only single tilts in which the pattern was clearly visible in the raw data were used. Sections of the spore envelopes were then selected using the “helical filament” picker, by manually clicking on the start and end points of the desired section. Particles were extracted every 40 Å using a box size of 470 Å. The results shown in the manuscript were obtained from 1,726 particles after two rounds of 2D classification and removal of duplicates.

### Subtomogram averaging of the cell envelope

For subtomogram averaging of the cell envelope, the best results were obtained using tomogram 108 from dataset 2. Particles were picked in dynamo^86^ using the membrane models where particles are regularly distributed on a selected surface and extracted such that the orientation of the subtomogram relative to the membrane can be used as a prior. A first round of refinement was then performed with a limited cone aperture to prevent flipping, and to make sure that the subtomogram orientation remained perpendicular to the membrane during alignment. The shift limits, cone and azimuth sampling were reduced progressively during the 15 iterations of refinement. Duplicate particles were removed in the last round. This led to a thick outermost layer followed by a bi-layer. A second round of refinement, using a mask on the exospore led to the most resolved pattern. Using the PlaceObject tool^87^, we observed the correlation of each subtomogram with the consensus model, and saw a large patch at the center of the selected envelope section with higher correlation.

### Subtomogram averaging of polar tubes

10 well-contrasted tomograms showing clear and long enough portions of the PT sections were selected for subtomogram averaging (STA), and the PT sections from each cell were processed independently. The overall workflow is presented in **Extended Data Fig. 7** and was inspired by the workflow in Laughlin et al.^88^. Briefly, in step 1, we first performed global STA to refine coil sections covering the whole diameter of the tubes. Each set of subtomograms (STs) was then aligned globally. In step 2, the aligned centers were used to pick new subtomogram averages centered at different radii from the center to refine the average of features located either on the OF-layer or on IF-layer. *Step 1*. Subtomograms were first picked manually using the filament (“crop along axis”) model in dynamo^86^ run within Matlab (R2019b, Natick, Massachusetts: The MathWorks Inc.), using separation of 3 to 5 pixels. To ensure the orientation of the different ST were coherent, these were picked from the back to the front of the tomogram for the coils on one side of the spore, and from the front to the back for the coils on the opposite side of the spore. An initial model was obtained by adding all manually picked STs together prior to any refinement, and radial pre-filtering was done by applying a high circular symmetry (C57). The rough cylinder obtained was used as a reference for the first round of refinement. Refinement was then performed in dynamo-v1.1.514^86^, during which the out-of-plane searches were restricted to prevent ST from flipping sidedness, and duplicates were removed in the last iterations. The intermediate averages contain 183 to 1,055 subtomograms depending on the spore (**Supplementary Table 3**), with reported resolution estimated to be below 50 Å (FSC 0.5 after splitting the data in dynamo). To estimate the radii of each layer from each STA, we computed 2D radial averages within Matlab using home-made scripts. The position of the different layers (M-layer, OF-layer, the IF-layer and the C-region) were then estimated by looking at local peaks on the radial averages of each STA.

*Step 2*. Focused refinement was carried out on the OF-layer and IF-layer to improve resolution. Smaller regions were picked at appropriate radii from the center of pre-aligned coil sections from Step 1. For each spore, the processing was done on the four tomograms that displayed the clearest filaments after step 1. The subtomograms picked on each layer were first processed in Dynamo (step 2a), then in Relion 4.0 (step 2b), and the best one in Relion 3.1 (step 2c). The initial model for each reconstruction (step 2a) was obtained by averaging the extracted ST without alignment. Refinements were performed within dynamo, and the azimuthal tilt was limited to reduce the risk of having the ST rotate 180° and maintain consistent orientation among subtomograms. To improve the resolution, we optimized masks for different regions of the layers. At the end of the refinements, duplicate particles were removed. Inspection of particle positions, orientations and correlation was done using PlaceObjects in Chimera. We noticed that portions of the PT contained regions where orientations of the ST were locally consistent and had higher cross-correlation with the consensus STA, while in other regions, the correlation was lower and adjacent STs appeared to be in random orientations relative to each other. These results are consistent with visual observations and manual segmentation, in which distinct densities could be used to manually segment filament-like structures from the tomograms in some regions, while in other regions, discrete densities could not be clearly identified. The STs in the most disordered regions were filtered out by setting a threshold on the cross-correlation value to the final model, as estimated in the dynamo tables generated after each refinement. We further optimized cleaning strategies such as only keeping STs for which orientations were consistent with the neighbors: we assumed that neighboring ST should have a similar orientation. To do this, we wrote a matlab script to remove those STs where at least one Euler angle was too different from at least 2 from its 5 closest ST neighbors (that is if at least one of its Euler angles differs by more than 30 degrees with at least 3 out of 5 of its nearest neighbors). After cleaning our subset of STs, and prior to transferring data to Relion 4.0 for further processing, we removed: 1) STs with lower cross-correlation to the consensus STA, 2) STs for which orientation was inconsistent with most of their neighbors 3) duplicates which appeared due to our over-picking strategy.

In step 2b, the most promising averages and STs were transferred to Relion 4.0 ^85^ via dynamo2relion code^89^. One round of local 3D refinement was run to prevent the relative orientation of the STs from diverging too much. Then, different strategies were optimized to improve the resolution, including: varying local refinements with different restraints on the angle and translation, 3D classification jobs with 1 to 5 classes and varying restraints on alignment, Estep or T values, and various masks on each layer, as well as on 1 to 4 filaments. In step 2c, one of the reconstructions was more featureful, displaying densities that may resemble repeating subunits. This dataset was transferred to Relion 3.1 after re-extraction of the subtomograms within WARP, and another round of 3D refinement was performed in Relion 3.1. Note that combining datasets from different spores did not improve the results. The best results were obtained from performing subtomogram averaging within a single cell, possible due to cell-to-cell heterogeneity. The final estimated resolutions of our models are described in **Supplementary Tables 4 and 5**.

Steps 2a and 2b were also applied to STs extracted at radii consistent with the position of IF-layer for the four tomograms in which that region showed the best contrast on the tomograms.

The orientation of the tilt axis used was provided by NCITU and NYSBC, where the tilt series were acquired, and defined using control samples. To verify that this information indeed led to the proper biological handedness in our reconstructions, we used prior information from the results of a previous study, which showed the PT is coiled within the spore as a right-handed helix^27^. One of our tomograms (from tomogram 1) contains several coil cross-sections as well as one crucial side-view highlighting the connection between left and right cross-sections, which allowed us to verify that this particular PT was indeed right-handed within the spore as expected.

### Measurements from cryo-ET data

#### Endospore thickness measurements

Endospore thickness was measured from 8 times binned, SART reconstructed tomograms in IMOD^78^. The slicer tool was used to orient a section of the endospore so that it appeared to have visually uniform thickness and the distance was measured from the plasma membrane to the inner layer of the exospore as this is the region where the endospore lies. For each tomogram, 3 measurements in 3 different positions were done.

#### Polaroplast membrane spacing

Polaroplast membrane spacing from tomograms 3, 15, 16, 25, 50 was measured using IMOD^78^. Distance was recorded from the center of one membrane to the center of the next membrane.

#### Nuclear Envelope thickness

Nuclear Envelope thickness was measured using IMOD^78^ from tomogram IDs: 1, 2, 4, 5, 8, 15, 28, 43, 47, 51, 52, 62, 79 and 94 where the nuclear envelope was clearly visible. Distance was measured from the center of the outer nuclear membrane to the center of the inner nuclear membrane.

#### Mitosome size measurements

As mitosomes are randomly oriented in our data, we measured the length along the longest axis of the mitosome visible in the tomogram using IMOD^78^.

#### Distances between segmented filaments and putative subunits

Distances between consecutive filaments from manual segmentation and STA for the OF-layer were measured in 3D using chimera^84^ by placing an atom on consecutive filaments and measuring the center to center distance. The same approach was used to measure the distance between putative subunits. The angle of the filament in the OF-layer to the vertical axis was measured by creating two axes in chimera^84^: 1) A filament axis tangent to the filament 2) A corresponding vertical axis. The angle between these axes was measured. The focused STA showing filaments on both OF and IF-layer displayed an opposite handedness and a relative angle of about ∼50 degrees between them, and was used to infer the angle of the filaments in the IF-layer to the vertical axis of the PT. The distance between consecutive filaments in the IF-layer was measured the same way as for the OF-layer.

#### Determination of number of filaments in OF-layer using segmentation

For this analysis, 7 coil sections from tomogram 4, where the filament segmentation was clearest, were selected. Tomogram 4 was opened in IMOD^78^ and using the slicer tool, an image was taken for each coil by orienting the tomogram so that the coil appeared circular. These images were then imported into FIJI^81^ where the circumference of the OF-layer was measured by drawing a segmented line around the densities. To determine the number of filaments *N*, the circumference *circ* was divided by the spacing *dist_x* between filaments in the x axis:

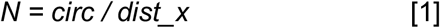

*dist_x* being calculated with:

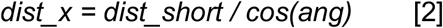

with *dist_short* the shortest distance between consecutive filaments (as described above), and *ang*, the angle between the filaments and the longitudinal axis of the tube, both measured from the corresponding segmentations.

#### Determination of filament number in OF-layer using STA

The filaments were counted manually in 3D in chimera^84^ by placing an ‘atom’ to track counts. This analysis was performed on the STA from the following tomograms: 1, 3, 4, 15, 32 and 84. The filaments were not clear enough to count all the way around in the following tomograms: 28, 41, 47 and 69 so these were excluded from the analysis.

#### Size of repeat unit in OF-layer

The most featureful map from focused STA was imported into Chimera^84^ and segmented to isolate putative subunits. Given the relatively low resolution, it is difficult to precisely identify the correct threshold and we therefore identified a lower and upper threshold which seemed to lead to acceptable map representations. The first unit reported volume of 61 Å^3^ to 94 Å^3^. The same process was repeated for a different subunit which reported volumes of 67 Å^3^ to 117 Å^3^. We took an average of the upper and lower bounds for the volumes as 64 Å^3^ and 105 Å^3^ and calculated the size in KDa assuming a globular protein occupancy around 1.21 Å^3 90^.

## Cellular modeling

All 3D models were built using Autodesk Maya software and exported as .obj files. MeshLab software was used to convert these into .stl files that could be inspected using UCSF Chimera.

The .stl files are available at github (https://github.com/mahrukhu/Polaroplast-Models). Six initial 3D models were generated to explore the range of potential organization of the polaroplast, with respect to their relation to the polar tube, and their folding pattern and axis (**Supplementary Table 6** and **Fig. 4A**). Each model was created with two variations, to accommodate variability in the length of the polaroplast. All other spore structures were modeled according to the SBF-SEM data from Antao et al^14^. The models were inspected using UCSF Chimera’s side view tool, and scored independently by 5 of the authors for their ability to correctly recapitulate the polaroplast organization in 11 different tomograms, representative of different section orientations. The individual scores were averaged to obtain a score for each sample/model pair, which were themselves averaged to obtain a final score for each model. Guidelines for scoring are described in **Supplementary Table 7**. Detailed scores are available in **Supplementary Table 8**. The ultrastructure of the polar tube was modeled using the experimentally-determined parameters listed in **Supplementary Table 9**.

