## Extended Data Figures, Supplementary Movie Legends and Supplementary Tables 1,3-9 for "Cryo-ET reveals the *in situ* architecture of the polar tube invasion apparatus from microsporidian parasites"

#### Extended Data Figure 1

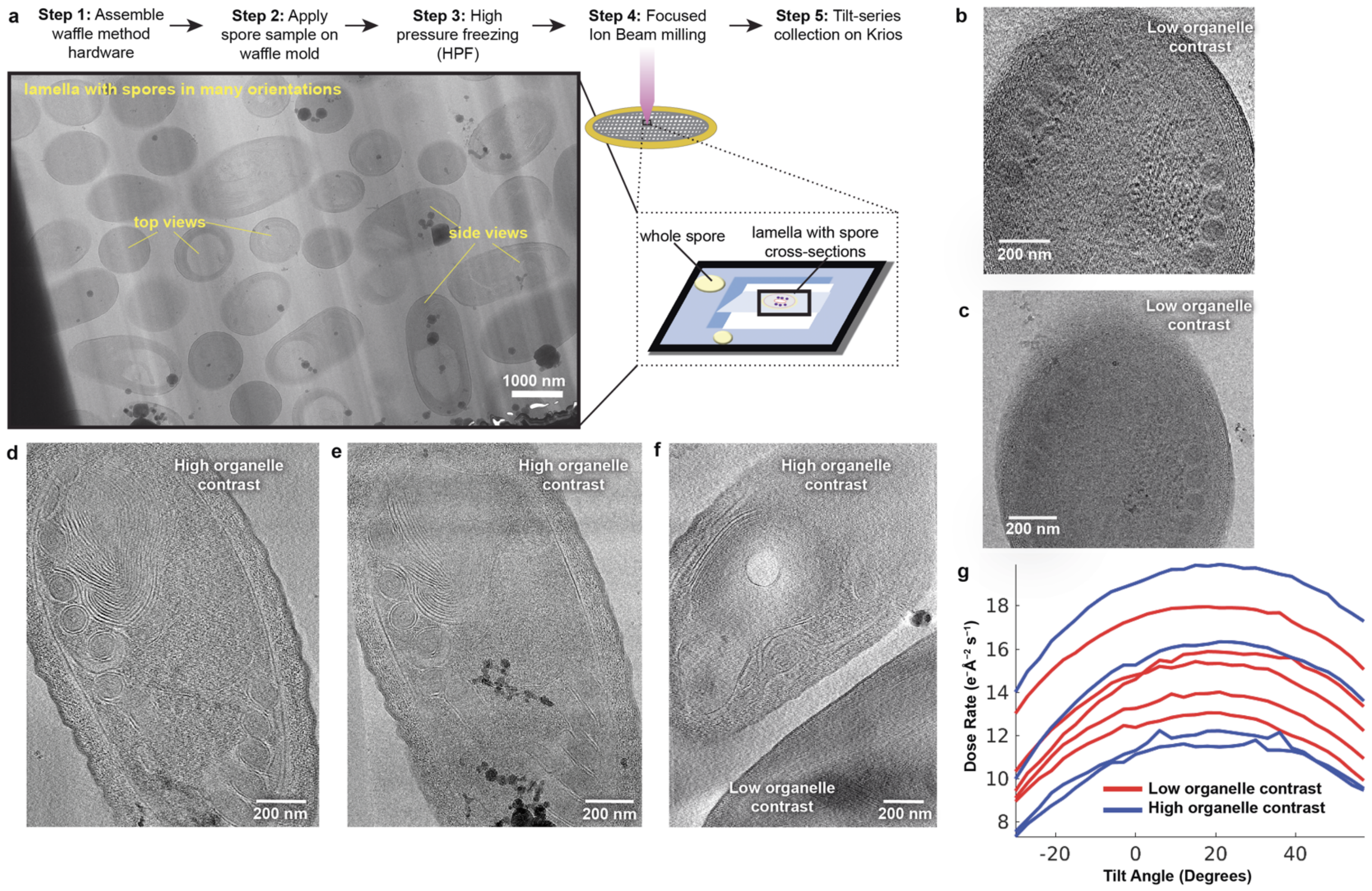


**Extended Data Figure 1. Data collection and cell-to-cell heterogeneity. (a)** Workflow for freezing spores, cryo-FIB milling and data collection. **(b)** A slice through a tomogram in which the organelles and membranes are not clearly distinguishable against the spore cytoplasm. **(c)** An image from the tilt series corresponding to the spore in (b). **(d)** A slice through a tomogram of a spore in which organelles and membranes are clearly defined with high contrast. **(e)** An image from the tilt series corresponding to the spore in (d). **(f)** Two spores captured in the same tilt series, one with low organelle contrast and one with high organelle contrast **(g)** Dose rate used as a proxy for ice thickness. There is no correlation between ice thickness and the observable organelle contrast in different tilt series.

#### Extended Data Figure 2

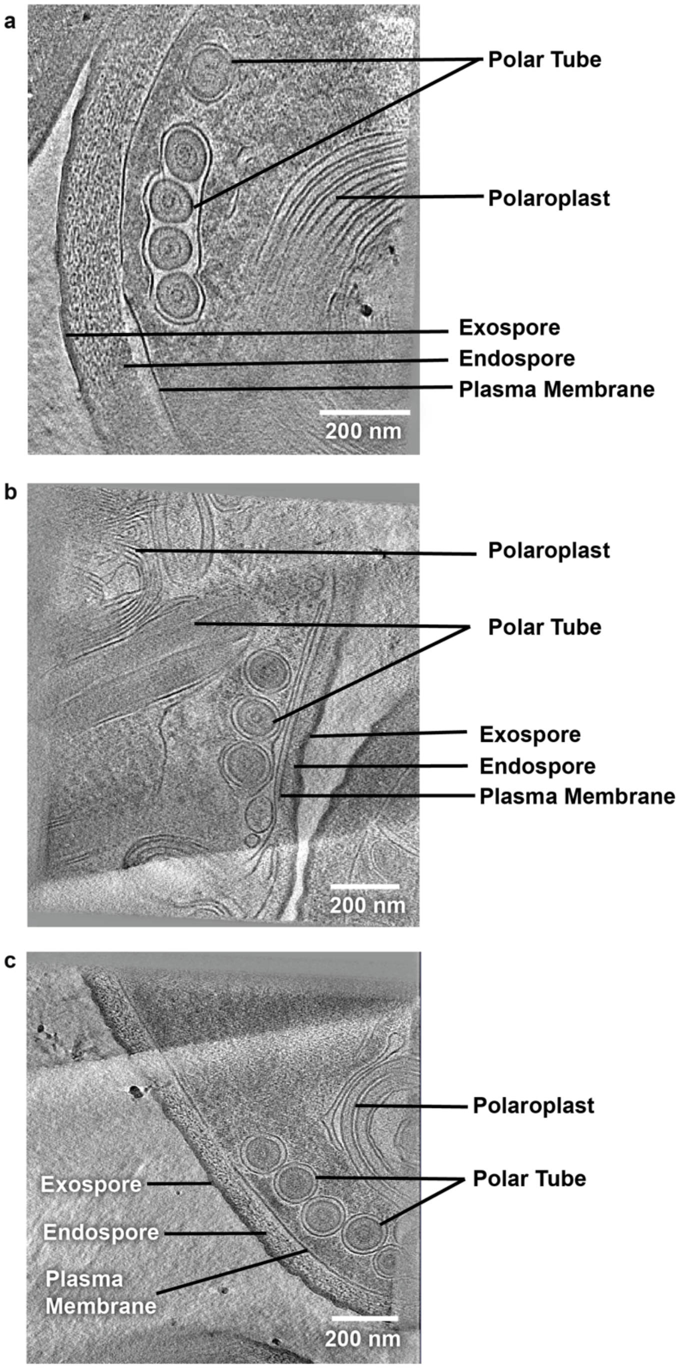


**Extended Data Figure 2. Representative images of dormant *E. hellem* spores.** 2D slices through tomograms of dormant *E. hellem* spores highlighting key cellular features including the PT, polaroplast and spore cell envelope. Three examples are shown in **(a)**, **(b)** and **(c)**.

**Extended Data Figure 3**

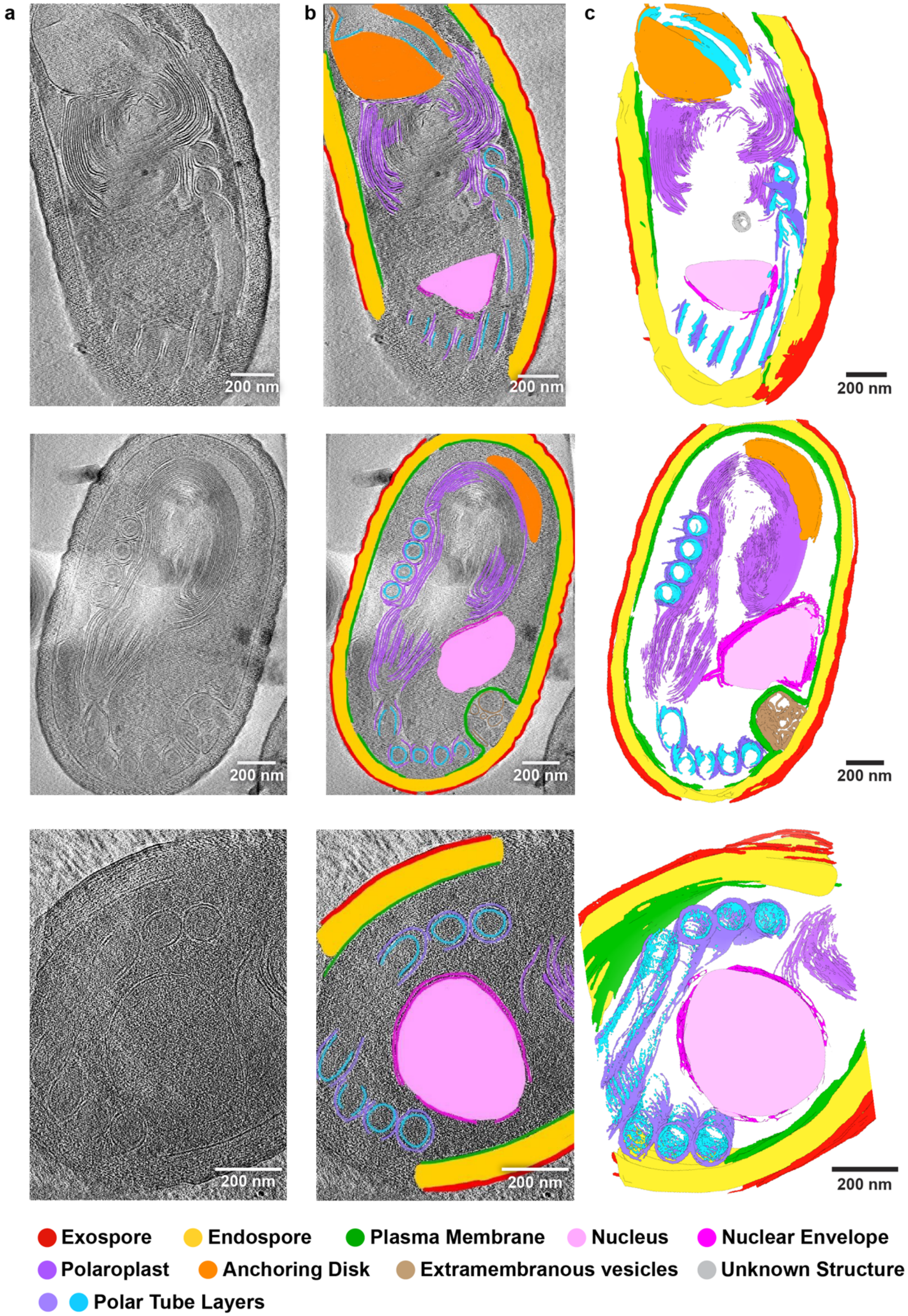


**Extended Data Figure 3. 3D segmentation of dormant *E. intestinalis* spores. (a)** 2D tomogram slices through spores. **(b)** Organelles annotated on tomogram slices corresponding to (a). **(c)** 3D reconstructions of the spores shown in (a) and (b).

#### Extended Data Figure 4

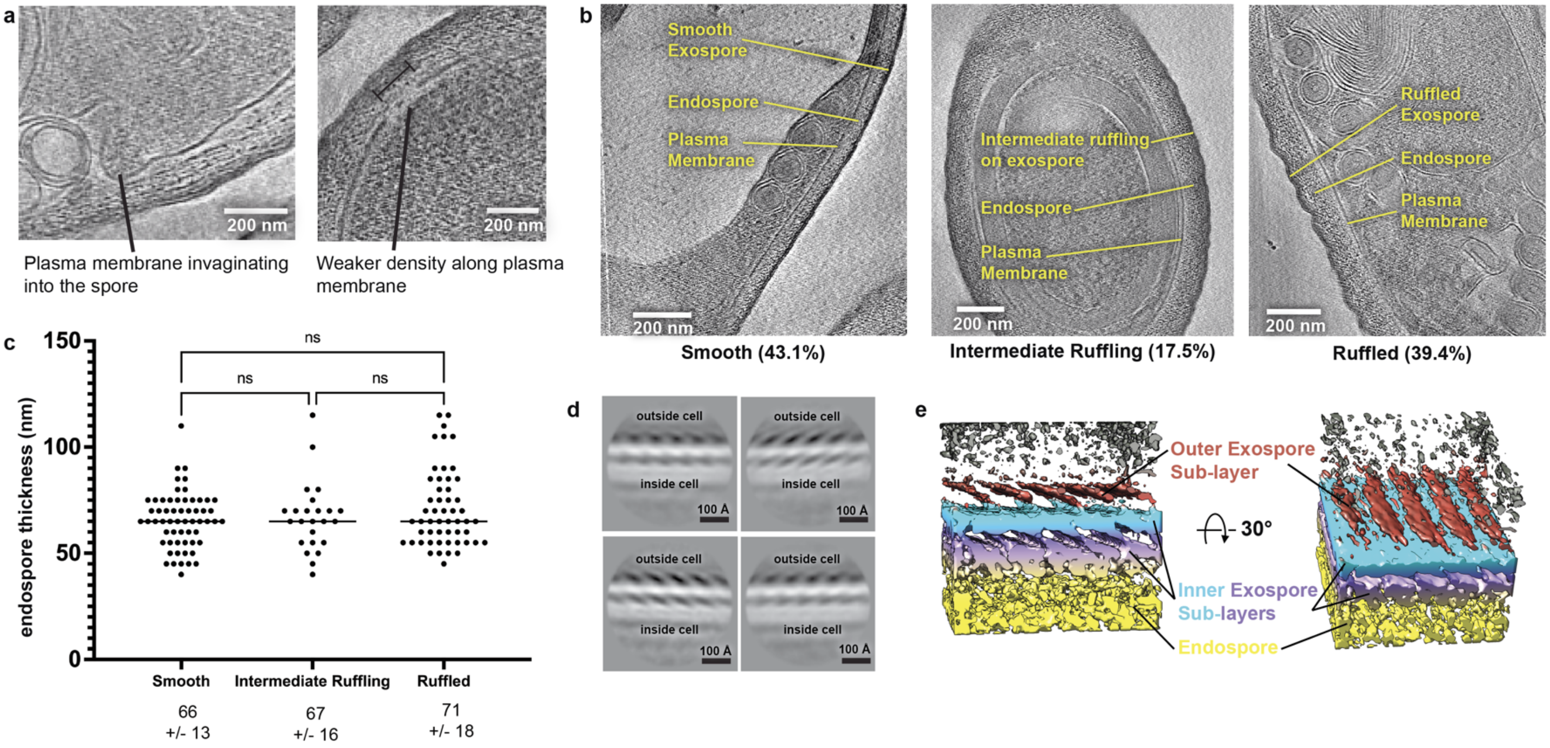


**Extended Data Figure 4. Organization of the spore cell envelope. (a)** Examples of plasma membrane features observed, including invaginations and regions with weak density. **(b)** Examples of spores that show different degrees of spore coat ruffling. **(c)** Graph showing quantification of endospore thickness as it is related to ruffling of the spore coat. The endospore thickness is 66 nm (st. dev. = 13 nm) for spores with a smooth coat, 67 nm (st. dev. = 16 nm) for spores with a slightly ruffled coat, and 71 nm (st. dev. = 18 nm) for spores with a ruffled coat. **(d)** 2D classification of the exospore, showing the presence of a repetitive pattern. **(e)** Subtomogram averages of exospore, gaussian filtered 0.5 Å, This is the same as Fig. 2D, but without dust hidden in Chimera.

**Extended Data Figure 5**

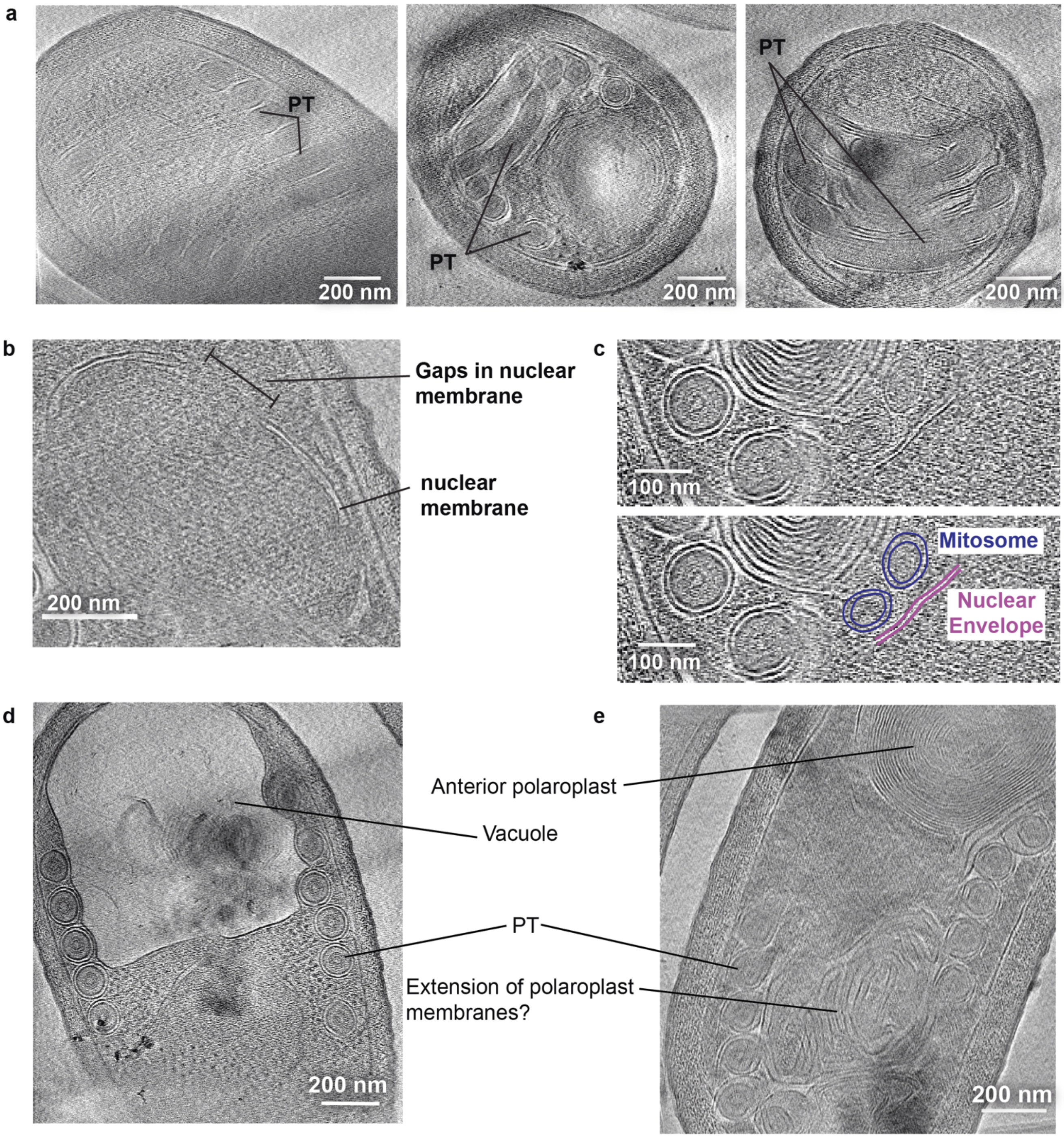


**Extended Data Figure 5. Organelle features observed in the dormant spores (a)** Examples of the PT observed in different orientations. **(b)** A slice through a tomogram showing a spore nucleus, with gaps in the nuclear envelope that may correspond to nuclear pore complexes. **(c)** Tomogram slice, unannotated (top) and annotated (bottom). Two double-membraned organelles with diameters of ~74.5 nm and 87.5 nm are located close to the nucleus, which we assign as mitosomes. **(d)** A slice through the tomogram of a spore showing a large, electron-lucent structure which is likely the vacuole. **(e)** A slice through a tomogram of a spore showing a loose network of membranes at the anterior end of the spore which may be extensions of the polaroplast.

#### Extended Data Figure 6

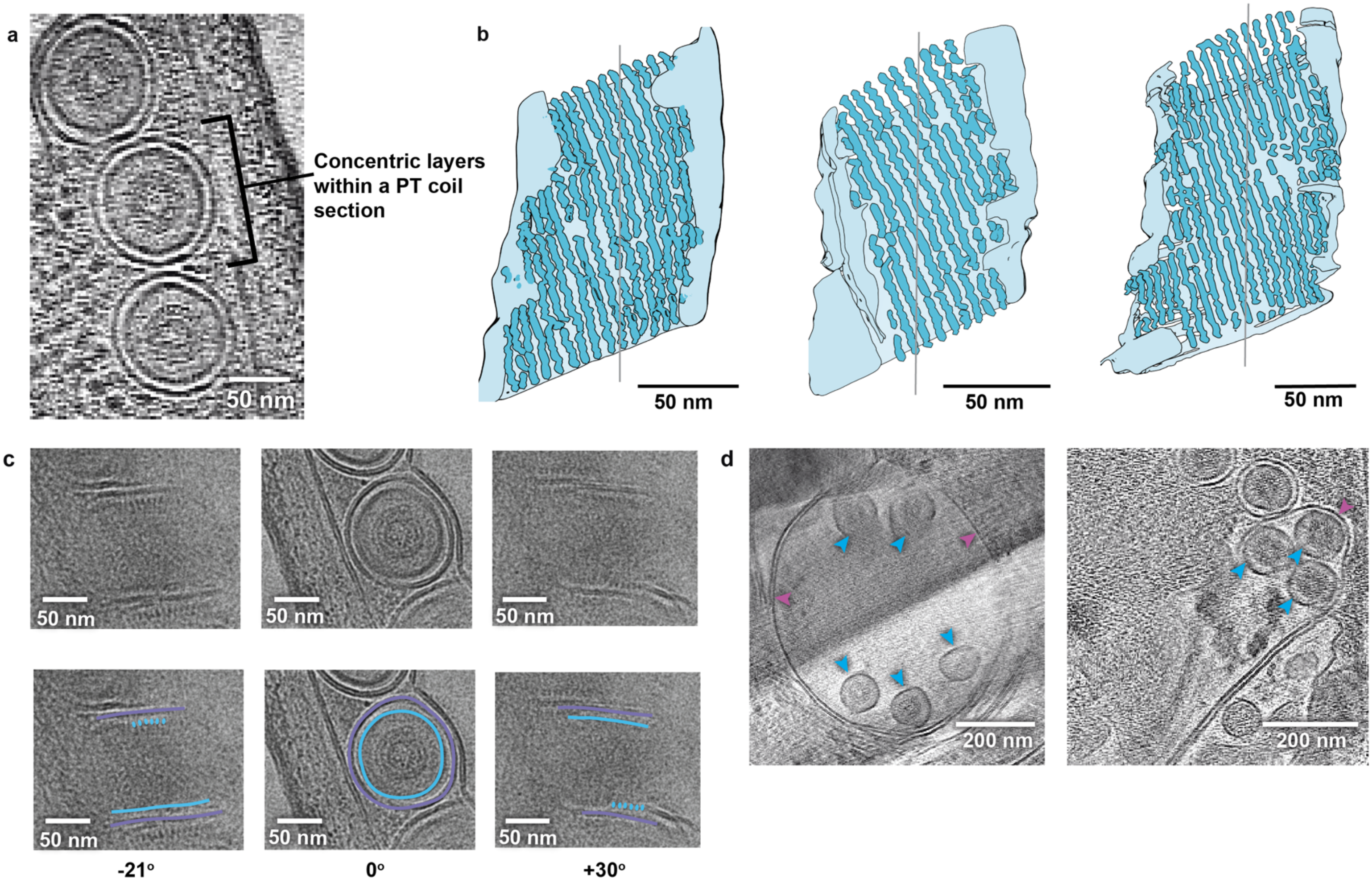


**Extended Data Figure 6. Features of PT morphology in dormant spores. (a)** Slice through a tomogram in which concentric layers of the PT are clearly visible. **(b)** 3D reconstructions of three PT coil segments, generated from segmentation of discrete densities in the OF-layer (see Fig. 3). Filaments are clearly visible upon 3D reconstruction of the segmentations. The spacing between these filaments is 66.3 Å to 73.8 Å and the angle of the filaments relative to the vertical axis is 22.6° to 24.3°. **(c)** A single PT coil section followed through a tilt series, unannotated (top) and annotated (bottom) to show M-layer (purple) and OF-Layer (cyan). At two opposing tilts, discrete densities on the OF-layer appear at opposite sides of the coil. Images are gaussian blurred using a sigma radius 2.00. **(d)** Examples of tomograms in which multiple PT coil sections (blue arrowheads) are observed within a single M-layer (magenta arrowheads).

#### Extended Data Figure 7

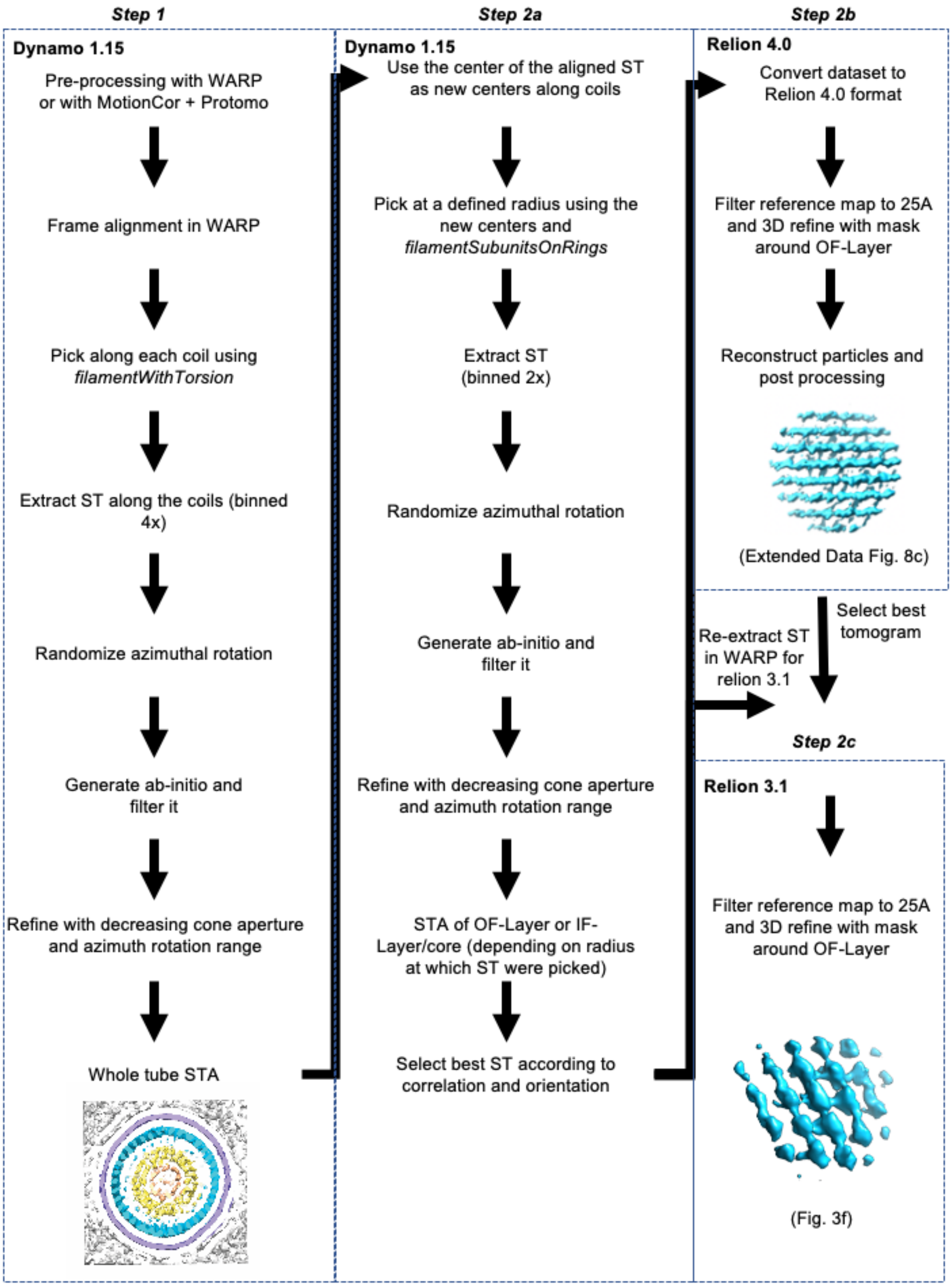


**Extended Data Figure 7. Workflow for subtomogram averaging**. The final workflow used to obtain subtomogram averages is presented here.

**Extended Data Figure 8**

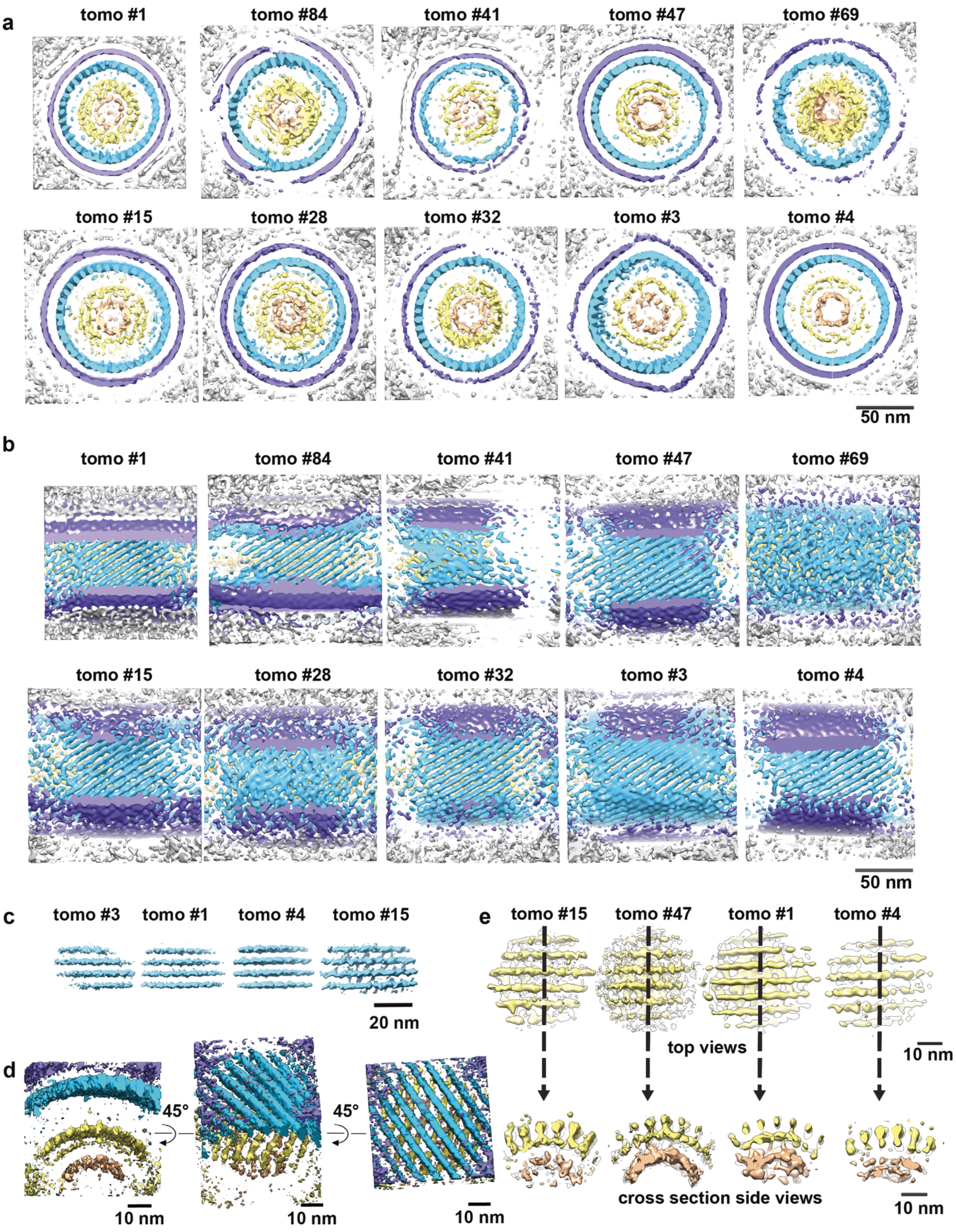


**Extended Data Figure 8. Additional data for subtomogram averaging. (a)** Front view cross sections from subtomogram averaging performed on whole sections of polar tube coils (10 spores from 3 different datasets).

**(b)** Side view cross sections of the same subtomogram averages as shown in panel A. **(c)** Subtomogram averaging of filaments in which the OF-layer was masked. **(d)** Same as Fig. 3e, prior to filtering and noise removal. **(e)** Subtomogram averaging of filaments in which the IF-layer was masked. Performed on 4 spores from 3 different datasets.

**Extended Data Figure 9**

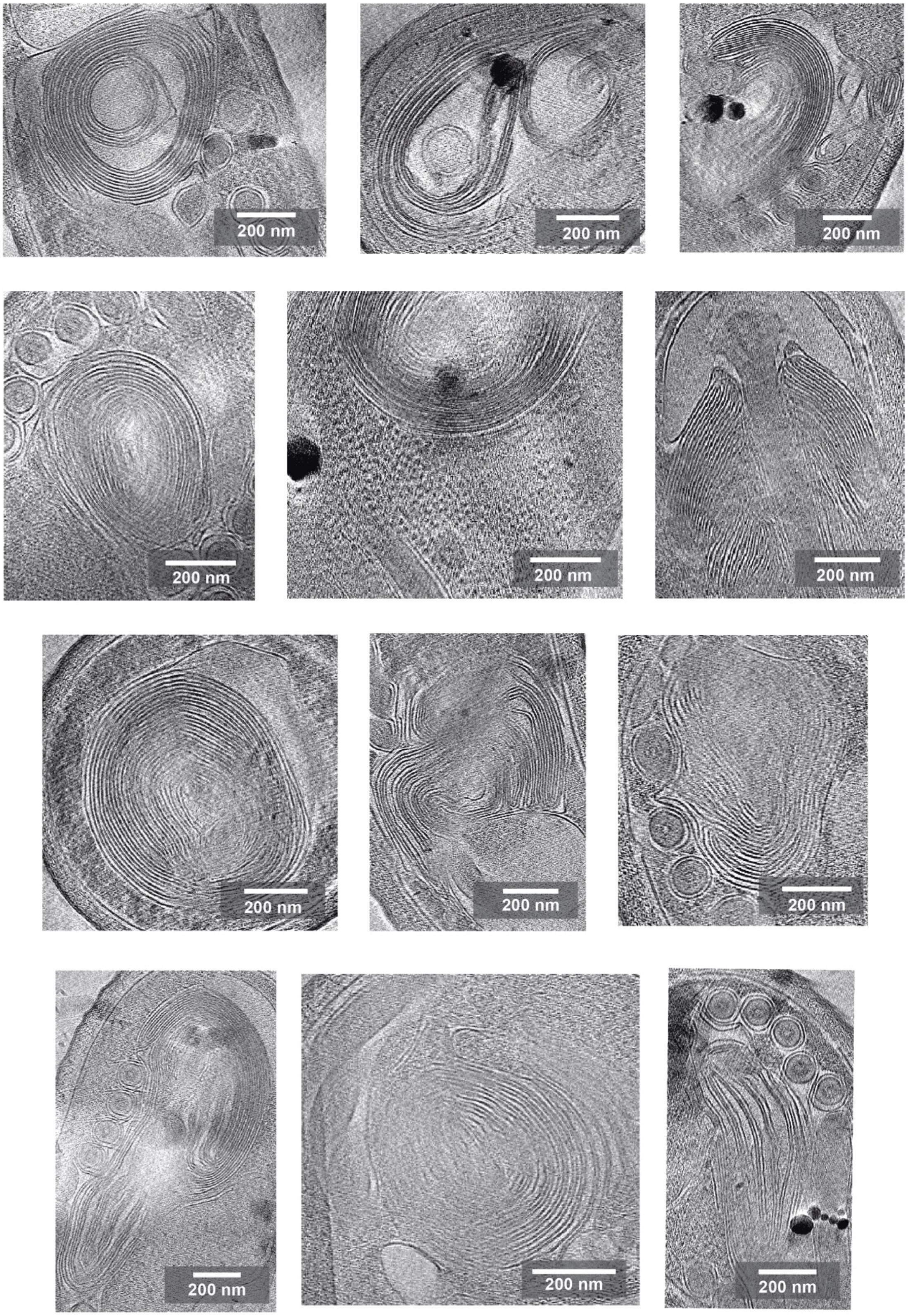


**Extended Data Figure 9. Polaroplast morphology observed in tomograms.** A gallery showing the different morphologies of the polaroplast observed across different spore tomograms, which is largely dependent on the angle of viewing.

### Extended Data Figure 10

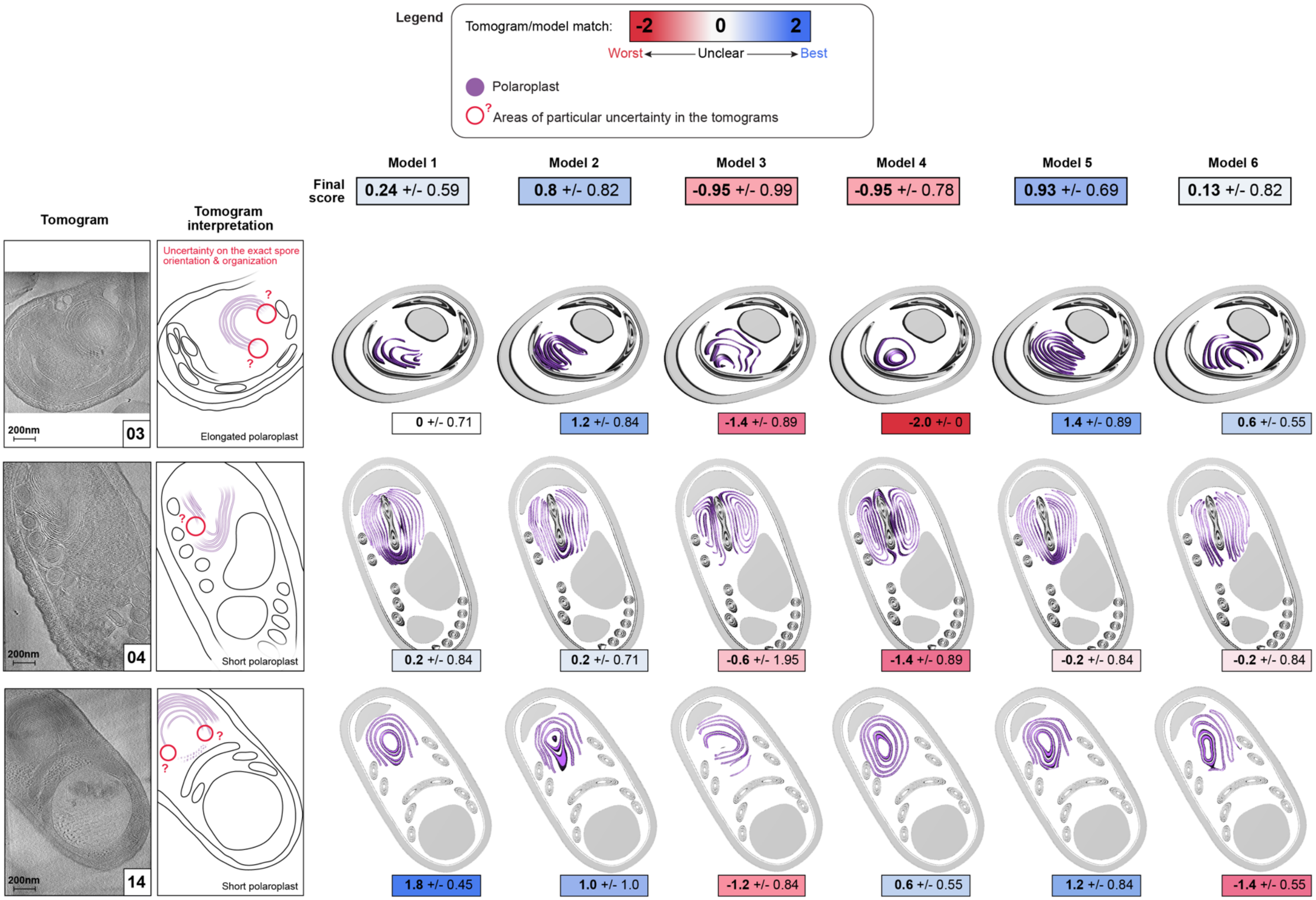


**(Figure continued on next page)**

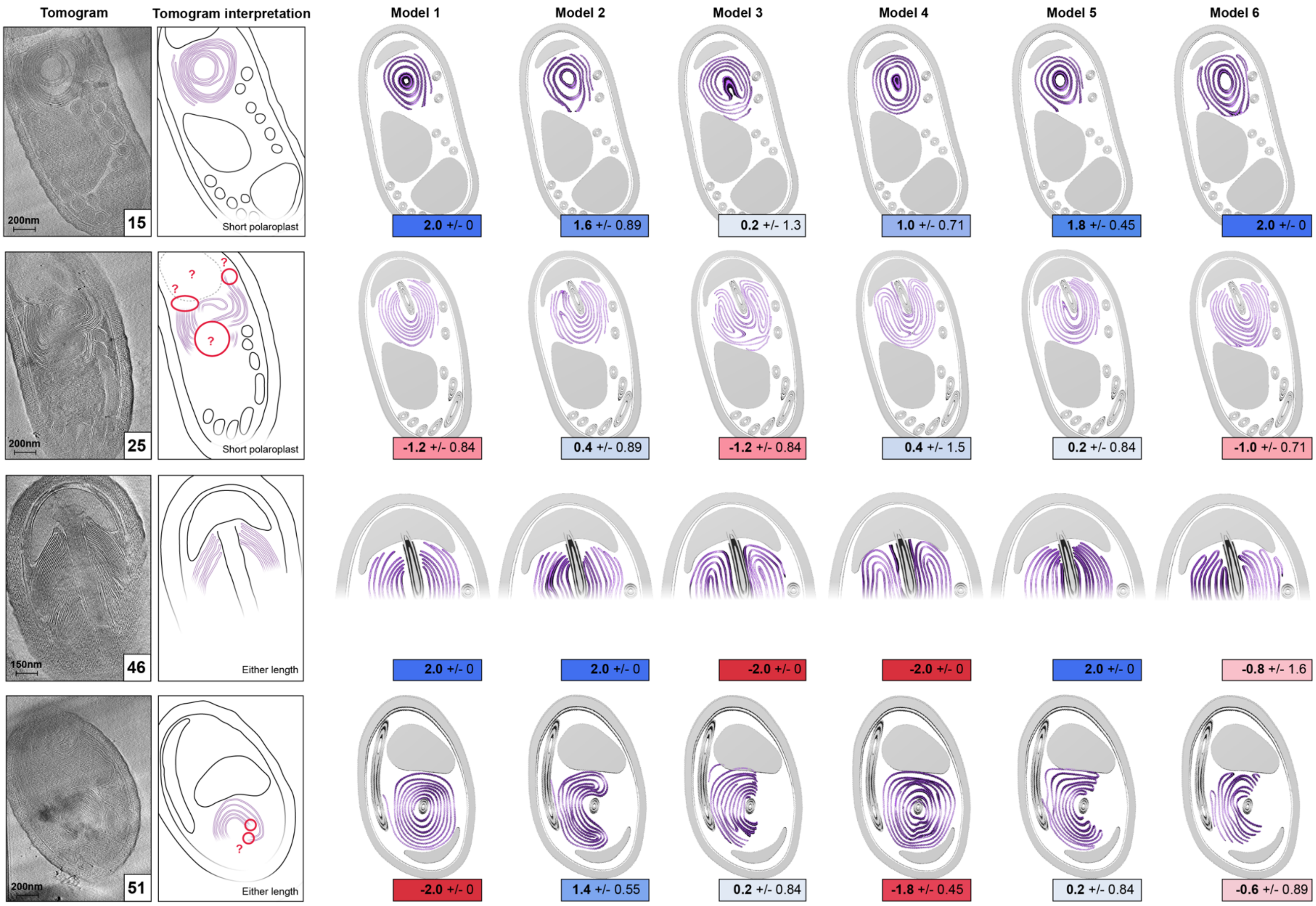


**(Figure continued on next page)**

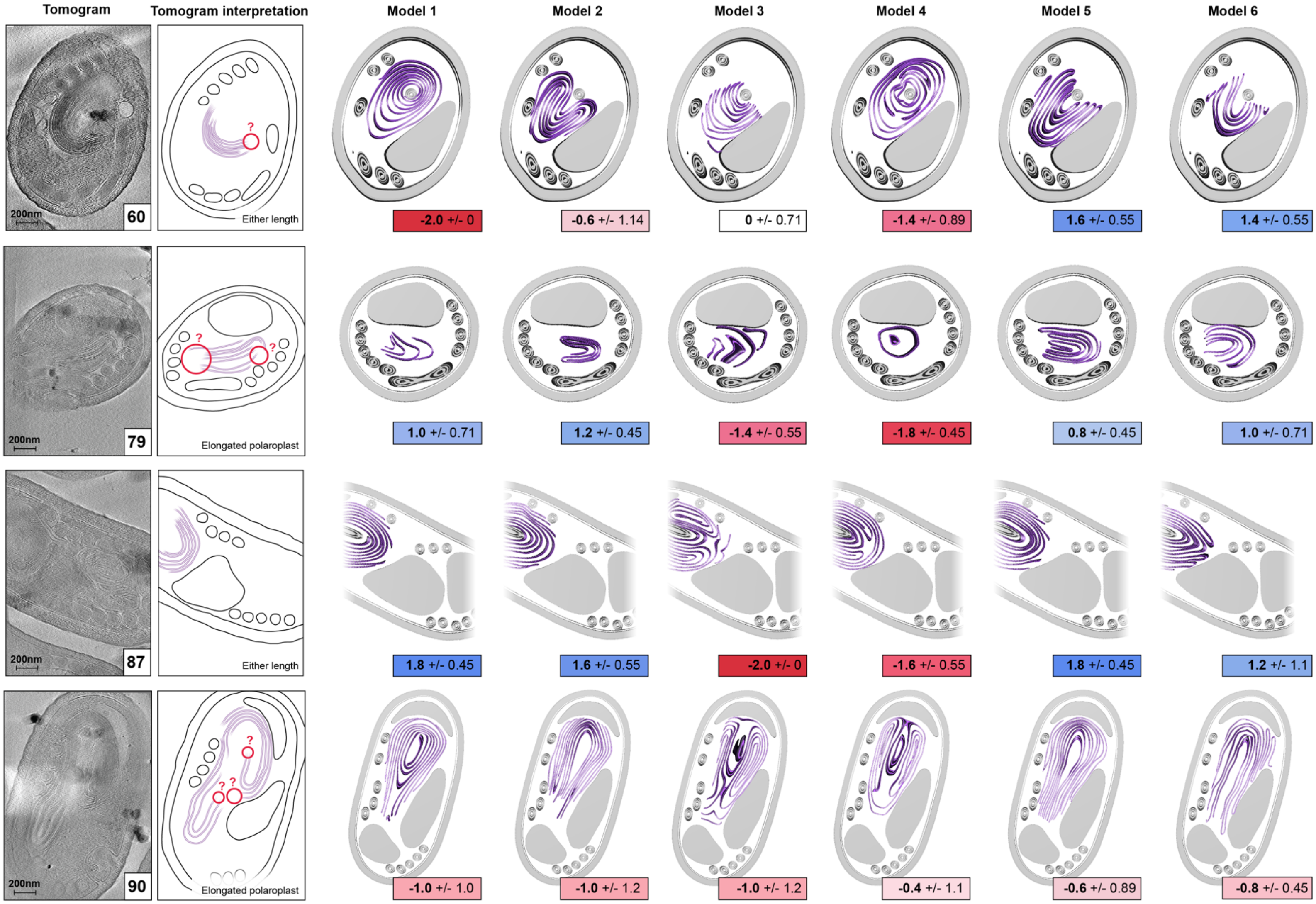


**Extended Data Figure 10. Virtual polaroplast models generated from cellular modeling compared with a set of tomograms.** Virtual models 1-6 were manually cropped in chimera in a variety of orientations to best recapitulate the key cellular features observed in each tomogram shown. Slices through each tomogram were compared with cropped models. Each model-tomogram pair was then assigned a score ranging from -2 (worst match) to 2 (best match) blindly and independently by five authors, according to guidelines detailed in the methods section.

#### Supplementary Movie Legends:

**Supplementary Movie 1. 3D reconstruction of a segmented E. intestinalis spore from tomogram 15.** Related to Fig. 2b.

**Supplementary Movie 2. 3D reconstruction of a segmented E. intestinalis spore from tomogram 46.** The linear portion of the polar tube surrounded by polaroplast and tucked into the anchoring disk is clearly visible. Related to Fig. 2f.

**Supplementary Movie 3. 3D reconstruction of a segmented E. intestinalis spore from tomogram 25.** Related to Extended Data Fig. 3.

**Supplementary Movie 4. 3D reconstruction of a segmented E. intestinalis spore from tomogram 90.** Related to Extended Data Fig. 3.

**Supplementary Movie 5. 3D reconstruction of a segmented E. intestinalis spore from tomogram 1.** Related to Extended Data Fig. 3.

**Supplementary Movie 6. 3D reconstruction of a segmented E. hellem spore from tomogram 109.** This tomogram highlights the connection between the polaroplast and the M-layer of the PT. Related to Fig. 5a.

**Supplementary Movie 7. 3D reconstruction of a segmented E. intestinalis spore from tomogram 3.** This tomogram highlights the connection between the polaroplast and the plasma membrane. Related to Fig. 5b.

**Supplementary Movie 8 - Model of a dormant microsporidian spore.** This movie synthesizes parameters from cryo-ET data to generate a model that is as complete and scientifically accurate as possible for the dormant microsporidian spore and the PT.

### Supplementary Tables

##### Supplementary Table 1. Cryo-ET data collection parameters

| | Dataset 1 | Dataset 2 | Dataset 3 | Dataset 4 |
|---|---|---|---|---|
| **Sample** | *E. intestinalis* | *E. intestinalis* | *E. intestinalis* | *E. hellem* |
| **Preparation** | Double step purification | Double step purification | Single step purification | Single step purification |
| **Microscope** | Titan Krios | Titan Krios | Titan Krios | Titan Krios |
| **Voltage (kV)** | 300 | 300 | 300 | 300 |
| **Detector** | Gatan K3 camera | Gatan K3 camera | Gatan K3 camera | Gatan K2 camera |
| **Magnification** | 33000 | 26000 | 26000 | 42000 |
| **Pixel size (Å / pixel)** | 2.077 | 3.353 | 3.431 | 3.298 |
| **Acquisition mode** | counting | super resolution | counting | counting |
| **Nominal defocus (um)** | -7 | -6 | -5.5 to -4 | -6 |
| **Number of frames per tilt** | 12 | 12 | 10 | 15 to 23 |
| **Dose per tilt ($e^-/A^2$)** | 1.67 - 6.67 | 1.92 | 2.87 | 1.13 - 1.74 |
| **Exposure time per tilt (s)** | 0.3 to 0.6 | 0.6 | 0.998669 | 1.5 - 2.3 |
| **Acquisition scheme** | -60°/+60°, 3° step | -30°/+60°, 3° step | -30°/+60°, 3° step | -50°/+50°, 2° step |
| **Software** | leginon | Serial EM | Serial EM & PACE-tomo | Leginon |
| **Number of tilt series acquired** | 33 | 58 | 94 | 12 |
| **Number of tomograms used** | 19 | 37 | 81 | 9 |
| **EMPIAR ID** | EMPIAR-12176 | EMPIAR-12176 | EMPIAR-12177 | EMPIAR-12175 |
| **CZII portal ID** | Dataset 10436 | Dataset 10436 | Dataset 10437 | Dataset 10438 |

**Supplementary Table 2. Summary and notes for all analyzed tomograms. (see Excel spreadsheet)**

**Supplementary Table 3. Parameters calculated from subtomogram averaging of whole PT sections.**
Related to Supplementary Fig. 8a, b.

| Tomogram ID | Number of subtomograms | Diameter of M-layer (nm) | Diameter of OF-layer (nm) | Number of filaments: OF-layer | Diameter of IF-layer (nm) |
|---|---|---|---|---|---|
| **1** | 1,055 | 103 | 89 | 36 | 50 |
| **3** | 718 | 123 | 97 | 40 | 54 |
| **4** | 628 | 110 | 87 | 40 | 51 |
| **15** | 250 | 115 | 88 | 38 | 49 |
| **28** | 317 | 115 | 91 | N/A | 55 |
| **32** | 199 | 115 | 91 | 39 | 55 |
| **41** | 183 | 102 | 82 | N/A | 49 |
| **47** | 221 | 115 | 91 | N/A | 49 |
| **69** | 267 | 121 | 91 | N/A | 44 |
| **84** | 365 | 121 | 91 | 41 | 52 |

**Supplementary Table 4. Parameters from subtomogram averaging of the OF-layer.**

| **Tomogram ID** | 1 | 3 | 4 | 15 (Relion 3.1) | 15 (Relion 4.0) |
|---|---|---|---|---|---|
| **Number of sub-tomograms picked** | 19,149 | 36,232 | 9,366 | 7,353 | 7,353 |
| **Number of sub-tomograms in model** | 11,148 | 6,762 | 4,132 | 1,657 | 1,657 |
| **Reported resolution of unmasked volume from Relion (at FSC = 0.5 / at FSC = 0.143)** | 36 Å / 19 Å | 39 Å / 28 Å | 30 Å / 27 Å | 36 Å / 29 Å | 40 Å / 36 Å |

**Supplementary Table 5. Parameters from subtomogram averaging of the Core.**

| Tomogram ID | 1 | 4 | 15 | 47 |
|---|---|---|---|---|
| Number of sub-tomograms picked | 12,275 | 7,290 | 5,336 | 4,928 |
| Number of sub-tomograms in model | 4,405 | 1,194 | 4,033 | 3,180 |
| Reported resolution of unmasked volume from Relion (at FSC = 0.5 / at FSC = 0.143) | 38 Å / 31 Å | 45 Å / 41 Å | 46 Å / 20 Å | 36 Å / 29 Å |

**Supplementary Table 6**. **Criteria considered for cellular modeling of the polaroplast.**

| | **Model 1** | **Model 2** | **Model 3** | **Model 4** | **Model 5** | **Model 6** |
|---|---|---|---|---|---|---|
| **Relation to the PT** | PT enclosed | PT not enclosed | PT not enclosed | PT enclosed | PT not enclosed | PT not enclosed |
| **Folding axis relative to the long axis of the PT** | Parallel | Parallel | Orthogonal | Orthogonal | Parallel | Orthogonal |
| **Folding pattern** | Spiraling | Spiraling | Spiraling | Spiraling | Back-and-forth | Back-and-forth |

**Supplementary Table 7. Guidelines for scoring the six polaroplast models against cryo-ET data.** These guidelines were provided to each of five authors who blindly scored cryo-ET data against the six models.

| Score | Correspondence between the experimental image and the model |
|---|---|
| 2 | **Good match:**<br>There are no major uncertainties in the tomogram and the model recapitulates the polaroplast organization OR there is a hesitation between a finite number of ways to interpret the tomogram but the model would match anyway |
| 1 | **Potential match:** The tomogram present important areas of uncertainty but the model matches all the interpretable parts (or would with really obvious/minimal variation i.e. individual fold length or shape) |
| 0 | **Very unclear:**<br>- The model can't well recapitulate the orientation/subcellular organization of the spore seen in the tomogram<br>- OR the tomogram is too unclear to emit any judgment regarding the model |
| -1 | **Potential mismatch:** The model doesn't seem to match but the tomogram present some important uncertainties in key areas that prevent a definitive judgment |
| -2 | **Obvious mismatch:** The model clearly doesn't match the tomogram (regardless of potential uncertainties present in the tomogram) |

**Supplementary Table 8. Detailed breakdown of scoring of polaroplast models against cryo-ET data.** Scores from each of five authors for each spore are shown.

| Spore 03 | **Model** | **1** | **2** | **3** | **4** | **5** | **6** |
|---|---|---|---|---|---|---|---|
| | Score 1 | 0 | 2 | -2 | -2 | 2 | 1 |
| | Score 2 | 1 | 1 | -2 | -2 | 2 | 1 |
| | Score 3 | 0 | 2 | -2 | -2 | 2 | 1 |
| | Score 4 | -1 | 1 | -1 | -2 | 1 | 0 |
| | Score 5 | 0 | 0 | 0 | -2 | 0 | 0 |
| | **Mean** | **0** | **1.2** | **-1.4** | **-2** | **1.4** | **0.6** |
| | **StDev** | **0.71** | **0.84** | **0.89** | **0** | **0.89** | **0.55** |

| Spore 04 | **Model** | **1** | **2** | **3** | **4** | **5** | **6** |
|---|---|---|---|---|---|---|---|
| | Score 1 | 0 | 0 | -2 | -2 | 0 | -1 |
| | Score 2 | 0 | 0 | 2 | -2 | 0 | 0 |
| | Score 3 | 1 | 0 | -2 | -1 | -1 | 0 |
| | Score 4 | -1 | -1 | 1 | 0 | -1 | 1 |
| | Score 5 | 1 | 1 | -2 | -2 | 1 | -1 |
| | **Mean** | **0.2** | **0** | **-0.6** | **-1.4** | **-0.2** | **-0.2** |
| | **StDev** | **0.84** | **0.71** | **1.95** | **0.89** | **0.84** | **0.84** |

| Spore 14 | **Model** | **1** | **2** | **3** | **4** | **5** | **6** |
|---|---|---|---|---|---|---|---|
| | Score 1 | 2 | 2 | -1 | 0 | 2 | -1 |
| | Score 2 | 2 | 0 | -2 | 1 | 1 | -2 |
| | Score 3 | 2 | 0 | -2 | 0 | 0 | -1 |
| | Score 4 | 1 | 1 | 0 | 1 | 1 | -2 |
| | Score 5 | 2 | 2 | -1 | 1 | 2 | -1 |
| | **Mean** | **1.8** | **1** | **-1.2** | **0.6** | **1.2** | **-1.4** |
| | **StDev** | **0.45** | **1** | **0.84** | **0.55** | **0.84** | **0.55** |

| Spore 15 | Model | 1 | 2 | 3 | 4 | 5 | 6 |
|---|---|---|---|---|---|---|---|
| | Score 1 | 2 | 2 | 1 | 1 | 2 | 2 |
| | Score 2 | 2 | 0 | -2 | 2 | 1 | 2 |
| | Score 3 | 2 | 2 | 0 | 0 | 2 | 2 |
| | Score 4 | 2 | 2 | 1 | 1 | 2 | 2 |
| | Score 5 | 2 | 2 | 1 | 1 | 2 | 2 |
| | **Mean** | **2** | **1.6** | **0.2** | **1** | **1.8** | **2** |
| | **StDev** | **0** | **0.89** | **1.3** | **0.71** | **0.45** | **0** |

| Spore 25 | Model | 1 | 2 | 3 | 4 | 5 | 6 |
|---|---|---|---|---|---|---|---|
| | Score 1 | 0 | 1 | -1 | 0 | 1 | 0 |
| | Score 2 | -2 | 0 | 0 | 2 | 0 | -2 |
| | Score 3 | -1 | 1 | -1 | 1 | 1 | -1 |
| | Score 4 | -1 | 1 | -2 | 1 | 0 | -1 |
| | Score 5 | -2 | -1 | -2 | -2 | -1 | -1 |
| | **Mean** | **-1.2** | **0.4** | **-1.2** | **0.4** | **0.2** | **-1** |
| | **StDev** | **0.84** | **0.89** | **0.84** | **1.5** | **0.84** | **0.71** |

| Spore 46 | Model | 1 | 2 | 3 | 4 | 5 | 6 |
|---|---|---|---|---|---|---|---|
| | Score 1 | 2 | 2 | -2 | -2 | 2 | 2 |
| | Score 2 | 2 | 2 | -2 | -2 | 2 | -2 |
| | Score 3 | 2 | 2 | -2 | -2 | 2 | -2 |
| | Score 4 | 2 | 2 | -2 | -2 | 2 | -1 |
| | Score 5 | 2 | 2 | -2 | -2 | 2 | -1 |
| | **Mean** | **2** | **2** | **-2** | **-2** | **2** | **-0.8** |
| | **StDev** | **0** | **0** | **0** | **0** | **0** | **1.6** |

| Spore 51 | Model | 1 | 2 | 3 | 4 | 5 | 6 |
|---|---|---|---|---|---|---|---|
| | Score 1 | -2 | 1 | 1 | -2 | 0 | -1 |
| | Score 2 | -2 | 2 | 1 | -2 | 1 | -1 |
| | Score 3 | -2 | 2 | 0 | -2 | 0 | -1 |

| | | | | | | | |
|---|---|---|---|---|---|---|---|
| | Score 4 | -2 | 1 | 0 | -1 | -1 | 1 |
| | Score 5 | -2 | 1 | -1 | -2 | 1 | -1 |
| | **Mean** | **-2** | **1.4** | **0.2** | **-1.8** | **0.2** | **-0.6** |
| | **StDev** | **0** | **0.55** | **0.84** | **0.45** | **0.84** | **0.89** |

| Spore 60 | **Model** | **1** | **2** | **3** | **4** | **5** | **6** |
|---|---|---|---|---|---|---|---|
| | Score 1 | -2 | -1 | 0 | -2 | 2 | 2 |
| | Score 2 | -2 | 1 | 0 | 0 | 1 | 1 |
| | Score 3 | -2 | -2 | 0 | -2 | 2 | 2 |
| | Score 4 | -2 | 0 | -1 | -1 | 1 | 1 |
| | Score 5 | -2 | -1 | 1 | -2 | 2 | 1 |
| | **Mean** | **-2** | **-0.6** | **0** | **-1.4** | **1.6** | **1.4** |
| | **StDev** | **0** | **1.14** | **0.71** | **0.89** | **0.55** | **0.55** |

| Spore 79 | **Model** | **1** | **2** | **3** | **4** | **5** | **6** |
|---|---|---|---|---|---|---|---|
| | Score 1 | 1 | 1 | -2 | -2 | 1 | 1 |
| | Score 2 | 0 | 1 | -1 | -2 | 1 | 0 |
| | Score 3 | 2 | 2 | -2 | -2 | 0 | 2 |
| | Score 4 | 1 | 1 | -1 | -1 | 1 | 1 |
| | Score 5 | 1 | 1 | -1 | -2 | 1 | 1 |
| | **Mean** | **1** | **1.2** | **-1.4** | **-1.8** | **0.8** | **1** |
| | **StDev** | **0.71** | **0.45** | **0.55** | **0.45** | **0.45** | **0.71** |

| Spore 87 | **Model** | **1** | **2** | **3** | **4** | **5** | **6** |
|---|---|---|---|---|---|---|---|
| | Score 1 | 2 | 2 | -2 | -1 | 2 | 2 |
| | Score 2 | 1 | 1 | -2 | -2 | 1 | 0 |
| | Score 3 | 2 | 1 | -2 | -1 | 2 | 0 |
| | Score 4 | 2 | 2 | -2 | -2 | 2 | 2 |
| | Score 5 | 2 | 2 | -2 | -2 | 2 | 2 |
| | **Mean** | **1.8** | **1.6** | **-2** | **-1.6** | **1.8** | **1.2** |
| | **StDev** | **0.45** | **0.55** | **0** | **0.55** | **0.45** | **1.1** |

| Spore 90 | **Model** | **1** | **2** | **3** | **4** | **5** | **6** |
|---|---|---|---|---|---|---|---|
| | Score 1 | -1 | -2 | -1 | -1 | -1 | -1 |
| | Score 2 | -2 | 1 | 1 | 0 | -1 | 0 |
| | Score 3 | 0 | -1 | -2 | 0 | -1 | -1 |
| | Score 4 | 0 | -2 | -1 | 1 | 1 | -1 |
| | Score 5 | -2 | -1 | -2 | -2 | -1 | -1 |
| | **Mean** | **-1** | **-1** | **-1** | **-0.4** | **-0.6** | **-0.8** |
| | **StDev** | **1** | **1.22** | **1.22** | **1.14** | **0.89** | **0.45** |

| **Overall** | **Model** | **1** | **2** | **3** | **4** | **5** | **6** |
|---|---|---|---|---|---|---|---|
| | **Mean** | **0.25** | **0.80** | **-0.95** | **-0.95** | **0.93** | **0.13** |
| | **StDev** | **0.59** | **0.82** | **0.99** | **0.78** | **0.69** | **0.83** |

**Supplementary Table 9. Parameters for polar tube modeling.**

| **Parameter** | **Experimental value** |
|---|---|
| **Outer membrane layer** | |
| Diameter of layer | 114 nm (from STA) |
| **Outer filament layer** | |
| Filament spacing | 65.5 Å (from segmentation) |
| Angle to longitudinal axis | -25.3º (from segmentation) |
| Number of filaments | 38 |
| Handedness | left |
| Diameter of layer | 89 nm (from STA) |
| **Inner filament layer** | |
| Filament spacing | 60 Å (from STA) |
| Angle to longitudinal axis | +24.7º (from STA) |
| Number of filaments | 24 (from STA) |
| Handedness | right |
| Diameter of layer | 51 nm (from STA) |
| **Core** | |
| Diameter of layer | 39 nm (from STA) |
